# Plasmids promote bacterial evolution through a copy number-driven increase in mutation rate

**DOI:** 10.1101/2025.07.21.665944

**Authors:** Paula Ramiro-Martínez, Ignacio de Quinto, Laura Jaraba-Soto, Val F. Lanza, Cristina Herencias-Rodríguez, Adrián González Casanova, Rafael Peña-Miller, Jerónimo Rodríguez-Beltrán

## Abstract

Plasmids are autonomously replicating DNA molecules that stably coexist with chromosomes in bacterial cells. These genetic elements drive horizontal gene transfer and play a fundamental role in bacterial ecology and evolution. Theory suggests that plasmids might evolve faster than chromosomes, as the mutation rate per gene should proportionally increase with plasmid copy number. However, the segregation of plasmid copies to daughter cells is random, introducing an additional layer of genetic drift, known as segregational drift, that might delay plasmid evolution. The interplay between plasmid mutational supply and segregational drift determines the evolutionary rate of plasmid-encoded genes, yet the relative contribution of these opposite forces in plasmid evolution remains unclear. Here, we took a classical population genetics framework to devise a mathematical approximation that predicts the fate of plasmid mutations in bacterial populations. We then validate these predictions by integrating computational, experimental, and bioinformatic approaches. Our findings show that plasmid mutation rates scale logarithmically with copy number: while increasing copy number elevates the mutation rate, the effect diminishes at higher copy numbers, where additional copies yield only marginal increases. Nonetheless, the supply of new mutations consistently surpasses the impact of segregational drift across all copy number levels. These results underscore plasmids as powerful platforms for bacterial evolvability and help explain their remarkable prevalence across microbial phylogeny.

## Introduction

Since the origins of life, all species depend on genomic flexibility to adapt to the ever-changing environment. Life is contingent on permanent change. Bacteria readily acquire genomic changes through mutation and horizontal gene transfer (HGT). Mutation is inherent to DNA replication: no polymerase can copy DNA without mistakes. And it is from those mistakes that life evolved on Earth. However, excessive mutations cause organisms to lose their identity as gene function collapses and their fitness declines(Loewe and Hill 2010). It is thus not surprising that life balances evolvability with genome stability and has evolved mechanisms to keep mutations in check(Darmon and Leach 2014). But what if bacteria could benefit from specialized genetic platforms in which specific genes could evolve faster? Theoretically, this would be achieved by locally increasing the supply of mutations, either by directly increasing the rate at which mutations are produced (i.e., decreasing replication fidelity) or by increasing the target size of mutations (i.e., the number of DNA molecules that can acquire mutations)(Sano et al. 2014).

Plasmids —autonomously replicating DNA molecules that coexist with bacterial chromosomes— are ideal candidates to fulfil that role(Rodríguez-Beltrán et al. 2021). Plasmids are key agents of HGT and play a fundamental role in bacterial ecology and evolution, where they contribute to disseminating relevant traits such as antibiotic resistance(Wiedenbeck and Cohan 2011). Several factors highlight plasmids as potential drivers of bacterial evolution. Plasmid genes might evolve faster than chromosomal genes because they are not essential to their host (with notable exceptions, e.g., refs(Anda et al. 2015; Manzano-Marı’n et al. 2019) and are less connected to cellular metabolic and physiological networks(Cooper et al. 2010; Cohen et al. 2011; Tazzyman and Bonhoeffer 2015). This relieves plasmid-encoded genes of the functionality constraints typically associated with the evolution of chromosomal genes. Second, plasmids might be more prone to recombination than chromosomes(Niaudet et al. 1984; Rodríguez-Beltrán et al. 2015; Zhang et al. 2019). Recombination provides bacterial plasmids with extraordinary modularity, shuffling genes and combining beneficial alleles in the same backbone(Cooper 2007; Barlow et al. 2008). And third, plasmid-encoded genes experience an increase in gene dosage that typically translates into higher expression. This magnifies the selective impact of mutations with slight beneficial effects(Million-Weaver et al. 2012; Couce et al. 2015; San Millan et al. 2016).

Critically, most plasmids are stably present at more than one copy per cell (median= 3.02, IQR= 10.29 copies per chromosome; data from ref.(Ramiro-Martínez et al. 2025)), providing an effective polyploid genetic platform(Rodríguez-Beltrán et al. 2021). This raises complex evolutionary implications that are far from understood. On one hand, classical evolutionary theory predicts that the rate of new mutations should increase with gene copy number because each additional copy has an independent chance of acquiring a mutation. Therefore, due to their stable polyploid nature, any given gene should acquire mutations faster when encoded on a plasmid. Supporting this view, experimental evolution showed that *de novo* mutations conferring antibiotic resistance occur more frequently when the target gene is encoded on a plasmid(Martinez and Baquero 1990; San Millan et al. 2016). Moreover, numerical simulations and experimental approaches demonstrate that mutation rates per gene increase with plasmid copy number (PCN) as long as the mutations are dominant(Rodríguez-Beltrán et al. 2020; Santer and Uecker 2020a). This indicates that plasmids can enhance the rate of (beneficial) mutations, thereby increasing the evolvability of the genes they carry(Couce et al. 2015; San Millan et al. 2016; Rodríguez-Beltrán et al. 2020).

On the other hand, novel plasmid mutations experience complex allele dynamics (Rodriguez-Beltran et al. 2018; Rodríguez-Beltrán et al. 2021; Santer et al. 2022; Rossine et al. 2025)(**Figure 1a**). New plasmid mutations only appear in a single plasmid molecule. This mutated plasmid copy will co-exist intracellularly under heteroplasmy (also known as heterozygosity) with other non-mutated plasmid molecules until they eventually segregate into different cell lineages(Novick 1987; Rodriguez-Beltran et al. 2018). Since plasmid segregation is random, daughter cells inherit an allelic set that may differ from the one in the mother cell (for example, only wild-type copies). The fixation time of plasmid mutations is thus delayed by an additional layer of genetic drift termed segregational drift (Ilhan et al. 2019). Recent studies suggest that segregational drift hinders the evolution of plasmid-encoded beneficial alleles, as they are more prone to being lost from populations than chromosomal ones(Garoña et al. 2021; Garoña et al. 2023; Dewan and Uecker 2025; Pisera and Liu 2025).

**Figure 1.**
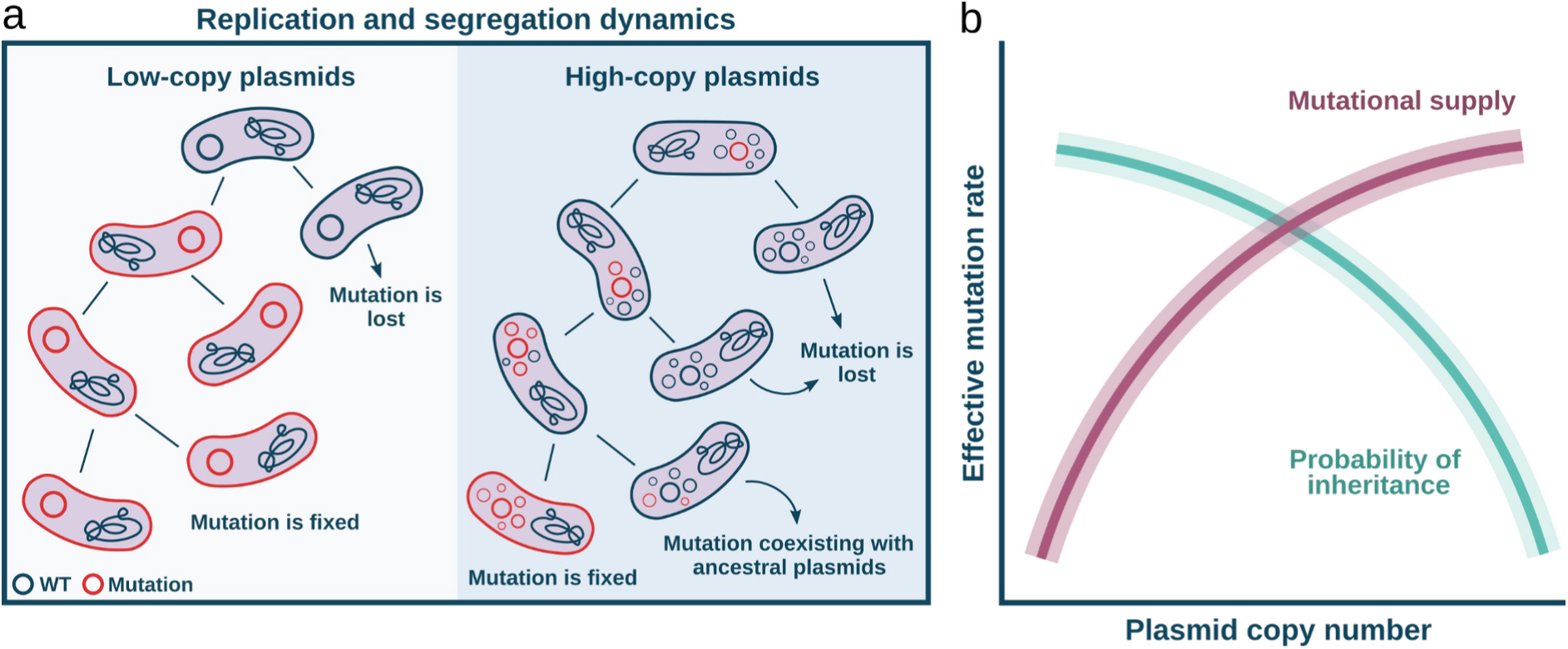
Plasmid-encoded genes face opposing evolutionary forces. **a)** Plasmid segregation dynamics. The left panel illustrates the segregation of plasmids with a single copy, where a mutant plasmid (red) is inherited during cell division. The right panel shows the dynamics of multicopy plasmids, where segregation occurs randomly upon division and mutant plasmids are either inherited or lost during cell division due to segregational drift. **b)** The illustration conceptually shows the trade-off between the probability of a new mutation arising (red line) and the probability of that mutation being inherited (green line) in subsequent generations. The mutational supply scales with PCN. In contrast, the chance that any given mutation is retained across generations decreases with PCN, as mutant plasmid copies are more likely to be lost due to segregational drift. This figure is illustrative and not based on real data.

Therefore, the evolutionary rate of plasmid-encoded genes depends on the interplay between two opposing forces: increased mutational supply and segregational drift (**Figure 1b**). However, the relative contribution of these two evolutionary forces remains controversial, with studies supporting(Couce et al. 2015; San Millan et al. 2016; Rodríguez-Beltrán et al. 2020) or opposing(Ilhan et al. 2019; Santer and Uecker 2020b; Garoña et al. 2021; Santer et al. 2022; Garoña et al. 2023; Dewan and Uecker 2025) the role of plasmids as potential catalysts of bacterial evolution. To settle this controversy, here we used a combined theoretical, computational, experimental, and bioinformatic approach to estimate how plasmid mutation rates scale with PCN.

## Results

### A plasmid-centered population genetics model predicts logarithmic increases in mutation rates as PCN increases

To investigate the mutational dynamics of plasmid-encoded genes, we first took a theoretical approach rooted in classical population genetics(Wakeley 2009; Etheridge 2011). Specifically, we developed a plasmid-centered population genetics model that shifts the focus from bacterial hosts to plasmids as independent evolutionary entities. In this approach, the key parameter N, representing population size in traditional population genetics, corresponds to the PCN. Thus, each bacterial reproduction event along the ancestral lineage is treated as a separate plasmid generation with a fixed population size set equal to the PCN.

We model plasmid inheritance using a discrete-generation approach related to the Wright-Fisher model(Durrett 2008), where plasmids act as individuals that replicate and segregate randomly across generations. More formally, our model falls within the broader Cannings class of population genetics models, which allows for variable offspring distributions while maintaining a fixed population size. During each cellular reproduction event, a random subset of plasmids is inherited, and the remaining copies are replenished by replication, preserving a constant copy number. This setup captures the random segregational dynamics of plasmids and allows us to trace the ancestry of plasmids across time (**Figure 2a**). Assuming that a low-probability neutral mutation can occur at each plasmid replication event, the count of distinct mutations in a sample correlates with the length of the genealogy tree. Consequently, the model suggests that nonlinear effects influence the genealogy of plasmid samples, and at certain scales, these genealogies align with the Kingman Coalescent universality class (**Figure 2b**, see methods). Under these conditions, the probability of two plasmids sharing a common ancestor decreases with PCN, and the total branch length of the genealogy grows logarithmically with copy number (**Figure 2c**). As a result, the effective mutation rate (µ_eff_), defined as the average number of distinct mutations per generation per plasmid, is expected to scale with the logarithm of PCN: *µ_eff_* =*μ* (*c*_1_+ *c*_2_ ln( *PCN* )), where *μ* denotes the underlying mutation rate (in mutations per generation and base pair) and *c*_1_and *c*_2_ are constants. This relationship suggests that increasing PCN raises the mutational supply, yet the rate of increase diminishes at higher PCNs (**Figure 2d**).

**Figure 2.**
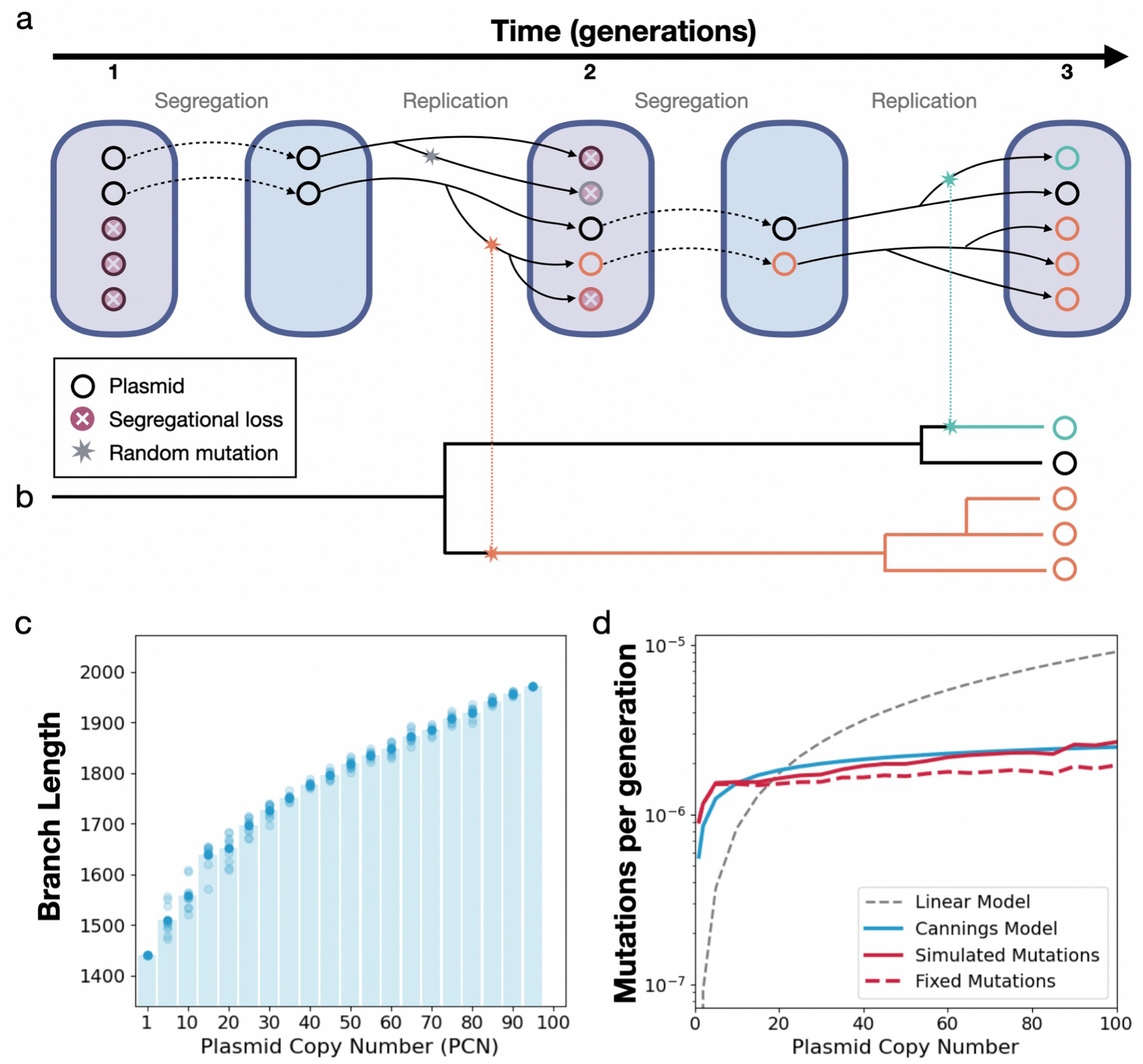
Population genetics model for plasmid mutation accumulation. **a)** Schematic diagram of the population genetics model. Each cell harbours a population of plasmids, with plasmid segregation during cell division modelled as a binomial process (p = 1/2). Reproduction events are represented within a Cannings framework, where PCN remains constant across generations. **b)** Example lineage tree of plasmids derived from the model. Genealogies demonstrate the coalescence dynamics driven by segregational drift, with branch lengths measured in generations, corresponding to the time to the most recent common ancestor. **c)** Total branch length of the lineage tree as a function of PCN. Branch length is defined as the cumulative distance from the root to the tips of the tree. Simulation results show that while total branch length increases with PCN, the rate of increase slows at higher PCN values. **d)** Number of mutations per generation per plasmid as a function of PCN. The gray dotted line represents a linear extrapolation based on the chromosomal mutation rate (_μ=9.2× 10_^−8^; see Figure 3), while the blue line depicts the Cannings model, where the constants c1 and c2 were adjusted to match the observed relationship between plasmid copy number and mutation accumulation. The red lines correspond to computer simulations: the solid red line shows the average total mutation rate across 1,000 simulations per PCN, and the red dashed line shows the average number of fixed mutations, defined as mutations present in all plasmids within a cell.

### Simulating multilevel evolution in multicopy plasmids

To evaluate the predictions of the population genetics model, we implemented a computational framework based on object-oriented programming principles (see methods). The model simulates bacterial populations as collections of individual cells, each containing plasmids modelled as independent entities (**Supplementary Figure 1**). This structure allows for the simulation of multi-level evolutionary processes, capturing key mechanisms — including random plasmid replication, segregation, and mutation— across multiple generations under controlled *in-silico* conditions.

In the model, each cell in the population contains a defined number of plasmid copies that replicate during cell growth until the target PCN is reached. Plasmid copies are randomly distributed to daughter cells during cell division, and a single bacterial cell is randomly selected at each generation to seed the next population, creating a bottleneck that drives genetic drift. Mutations occur with a non-zero probability in each plasmid copy during replication. Each mutation is recorded as an attribute of the corresponding plasmid object and includes details such as the generation of occurrence, allowing precise tracking of mutations over time within the simulation framework (**Supplementary Figures 2-5**).

We performed 1,000 independent simulations for a range of PCN, each elapsing 1,400 bacterial generations. At the end of the simulation, plasmids were sampled from the final bacterial population to reconstruct their evolutionary relationships. We then built a lineage tree by tracing the ancestry of each plasmid based on its generational history. Plasmids that share a common lineage (i.e., descend from the same ancestral plasmid in earlier generations) are grouped together, with branches representing points where lineages diverged due to random segregation events during cell division (**Supplementary Figure 6**).

Finally, we quantified the total length of the lineage tree, defined as the cumulative distance from the root to the tips across all plasmid lineages. Consistent with the theoretical predictions, as PCN increases and we sample more plasmids per cell, the total branch length also increases, reflecting more coalescing events and longer genealogical paths. However, as shown in **Figure 2c**, this increase slows at higher PCN levels, consistent with a logarithmic increase in tree length as the copy number rises, even though the total tree length continues to grow. Consequently, our computational model aligns with the theoretical prediction, and the number of mutations observed increases with PCN but at a diminishing rate (**Figure 2d; Supplementary Figure 7**).

### A mutation accumulation experiment validates computational and theoretical predictions

To experimentally test our theoretical and computational predictions, we used a mutation accumulation (MA) approach. MA experiments typically consist of serially passaging multiple replicate clonal lines by repeated streaking on rich agar plates to generate new colonies initiated by a single cell. This severe bottleneck minimizes natural selection, allowing genomes to accumulate nearly any mutation regardless of its selective effect, except those that are lethal or highly deleterious(Barrick and Lenski 2013). After a number of passages, mutations across entire genomes are identified by sequencing, allowing for direct estimation of mutation rates by simply counting the number of genetic changes accumulated over a defined number of generations(Halligan and Keightley 2009; Barrick and Lenski 2013).

To conduct the MA experiment, we used a hypermutator derivative (Δ*mutS*) of the *Escherichia coli* EPI300 strain carrying a plasmid with a tunable PCN. This plasmid, dubbed pTA44, can initiate replication from two distinct origins: *ori2* from the F plasmid and *oriV* from the RK2 plasmid(Aakvik et al. 2009). Replication from *ori2* is constitutive and leads to approximately one copy per cell. Replication from *oriV* depends on the presence of the replication initiation protein TrfA, whose expression is chromosomal in the EPI300 strain and controlled by the arabinose-inducible P_BAD_ promoter (**Figure 3a, Supplementary Figure 8**). Therefore, the PCN of the pTA44 plasmid shows a wide dynamic range spanning from ∼1 copy per cell in the absence of arabinose to nearly 70 copies at high arabinose concentrations (**Figure 3b**). Since bacterial growth was relatively robust to changes in PCN (**Supplementary Figure 9**), we selected three arabinose concentrations (0%, 0.0005%, and 0.05%) for subsequent experiments. These yielded, on average, 1, 10, and 60 plasmid copies per cell, corresponding to low, medium, and high PCN regimes, respectively (**Figure 3b**).

**Figure 3.**
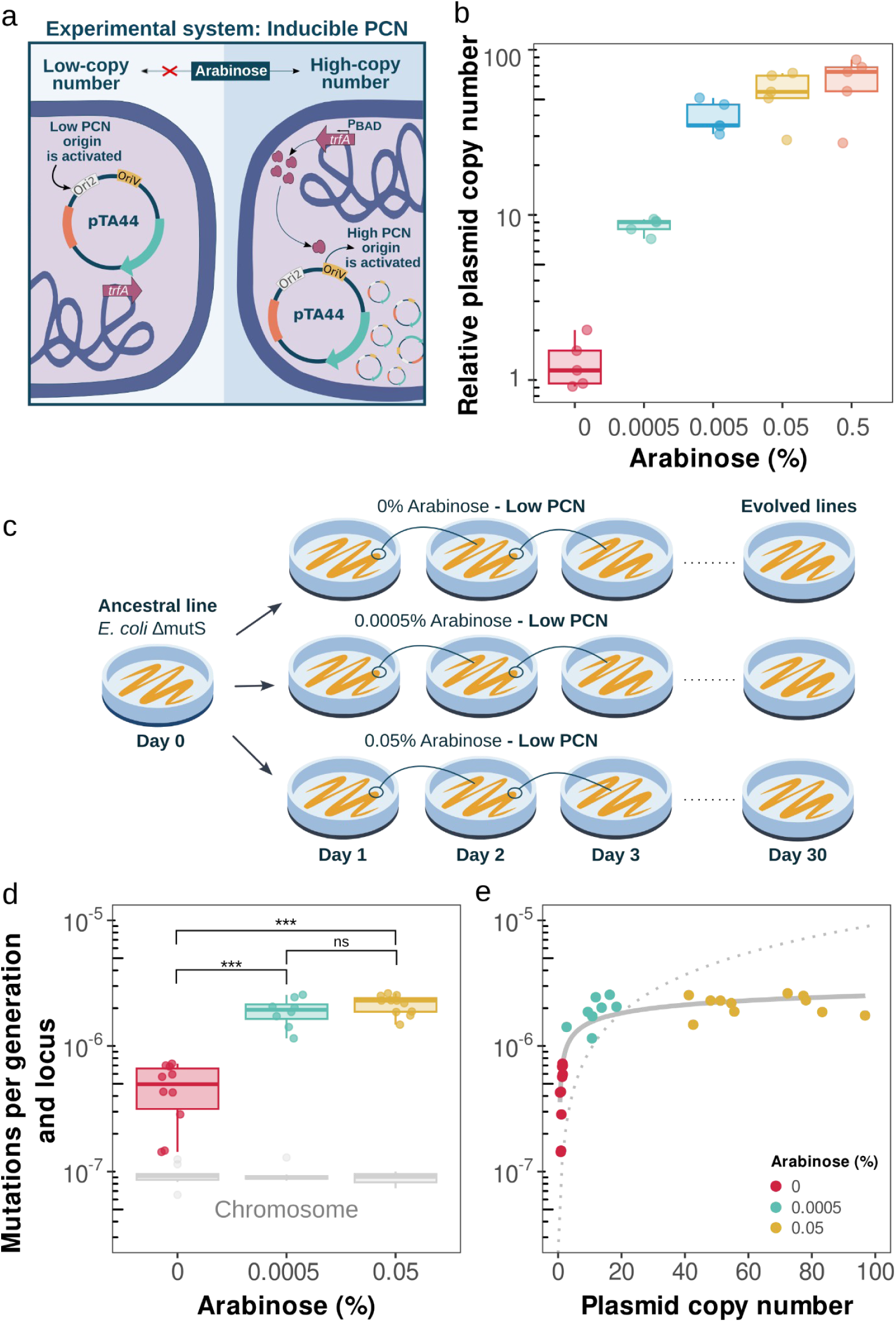
A MA approach validates theoretical predictions. **a)** Plasmid copy number induction in the pTA44 plasmid. In the presence of arabinose, expression of TrfA protein from the PBAD promoter is turned on, initiating replication from the high-copy replication origin (oriV). Without arabinose replication is driven by the low-copy replication origin (ori2). **b)** PCN of the pTA44 plasmid at different arabinose concentrations. The copy number of five biological replicates was calculated by qPCR. The average PCN of 9 technical replicates was 1.31 copies for 0% of arabinose, 9.32 for 0.0005%, 40.07 for 0.005%, 57.70 for 0.05%, and 67.65 for 0.5%. **c)** Experimental setup. We conducted a MA experiment at different arabinose concentrations. Each day, one random colony of each line was transferred to a new Petri dish with the appropriate arabinose concentration. This cycle was repeated for 30 days. **d**) Boxplots show plasmid mutation rates (mutations per generation and locus) at different arabinose concentrations, which correlate with plasmid copy number. Each point represents an independent replicate. Boxes indicate the median and interquartile range; whiskers extend to 1.5× IQR. Chromosomal mutation rates (grey) are shown for reference. Statistical significance was assessed using pairwise Wilcoxon rank-sum exact tests (*p < 10⁻⁴; ns, not significant). **e)** Plasmid mutation rates scale logarithmically with copy number (see **Supplementary dataset 2**). Each dot represents an independent experimental line, colored by the arabinose concentration used during evolution (see legend). The dashed line indicates the expected linear increase in the absence of segregational drift, based on the chromosomal mutation rate. The solid line shows the fitted Cannings model, with constants c₁ and c₂ adjusted to match the experimental data (see **Supplementary dataset 3**).

We serially propagated 12 independent lineages for each PCN regimen by daily restreaking a randomly selected colony onto fresh LB agar plates supplemented with the appropriate arabinose concentration over 30 days (**Figure 3c**). During each serial passage, bacterial populations expanded from a single cell to 10⁸ cells, corresponding to ∼24 bacterial generations (**Supplementary Figure 10**). This corresponds to approximately 670 generations per lineage and a total of ∼6,500 generations for each PCN regimen (**Supplementary dataset 1**). The number of generations was generally conserved across all lines throughout the experiment, suggesting that selection played a minimal role under our experimental conditions (**Supplementary Figure 10**, Spearman’s rank correlation test between number of generations and days, p = 0.81, rho = 0.014). Similarly, PCN was stable across arabinose concentrations, except in one line of the medium copy regimen (p5B3), which was excluded due to a *trfA* missense mutation causing a sharp decline in PCN (**Supplementary Figures 11–12**).

We sequenced the genomes of all evolved lineages and the ancestor and identified all mutations (single-nucleotide polymorphisms —SNPs—, and small indels). A key feature of plasmid segregation and replication dynamics is that multiple plasmid alleles can coexist within the same cell, leading to plasmid-mediated heteroplasmy or heterozygosity (Novick 1987; Rodriguez-Beltran et al. 2018; Garoña et al. 2021). Therefore, plasmid-encoded alleles can be polymorphic even in otherwise clonal lineages. To account for this possibility, we kept mutations with an allelic frequency of at least 5%, as done previously(Garoña et al. 2021). Following this approach, we identified 284 plasmidic and 7,620 chromosomal mutations (**Supplementary dataset 1**).

We calculated the spontaneous mutation rate for both plasmids and chromosomes by dividing the number of mutations by the length of each replicon and the number of generations elapsed for each line. As expected, chromosomal mutation rates remained unaffected by the presence of arabinose (Kruskal-Wallis rank sum test, p = 0.79; **Figure 3d**), and were consistent with previous estimations(Lee et al. 2012). In contrast, plasmid mutation rates were significantly higher in the medium and high PCN regimes than in the low PCN (Pairwise comparisons using Wilcoxon rank sum exact test, p < 10^−4^ between low and medium, and low and high copy number regimes; **Figure 3d**). There were no significant differences between the medium and high PCN regimes, despite a ∼5-fold difference in PCN (Wilcoxon rank sum test, p = 0.27). Strikingly, at the low PCN regime, where plasmids and chromosomes were present at comparable copy numbers (∼1 copy per cell), plasmid mutation rates were approximately five times higher than those of the chromosome.

We then analyzed the mutation rates of each evolved line individually. **Figure 3e** shows that plasmid mutation rates increased with PCN and were positively correlated (Spearman’s rank correlation, rho = 0.8, *p* < 10^−5^). However, the rate of increase diminished progressively as PCN rose. We observed a growing divergence between the observed mutation rate and the expected rate if mutations accumulated linearly with PCN, using the chromosomal mutation rate as a baseline (**Figure 3e**). This aligns with our theoretical and computational predictions, as a logarithmic regression best fitted the experimental data (R^2^=0.79; **Supplementary dataset 2**).

### Estimation of plasmid mutation rates from whole-genome sequencing data

We hypothesized that if mutation rates scale with PCN beyond experimental conditions, a compatible signature should be detectable in naturally occurring plasmids. To test this idea, we leveraged the DNA sequences available in the NCBI database.

In a typical whole-genome sequencing (WGS) experiment, a liquid culture is inoculated from a small number of cells and incubated overnight to allow bacterial growth. DNA is then extracted from the culture and sequenced. Raw sequencing reads are assembled into a consensus genome, representing the dominant genotype of the population. We reasoned that mutations could accumulate during the growth phase prior to DNA extraction in both chromosomal and plasmid DNA. While some of these mutations may be too infrequent to be detected, others may increase in frequency through genetic drift or selection and become detectable in the sequencing reads. Thus, aligning sequencing reads to their corresponding consensus assemblies would allow identifying within-sample mutations that likely arose during culture growth, and the number of such mutations would be roughly proportional to the underlying mutation rate (**Figure 4a**).

**Figure 4.**
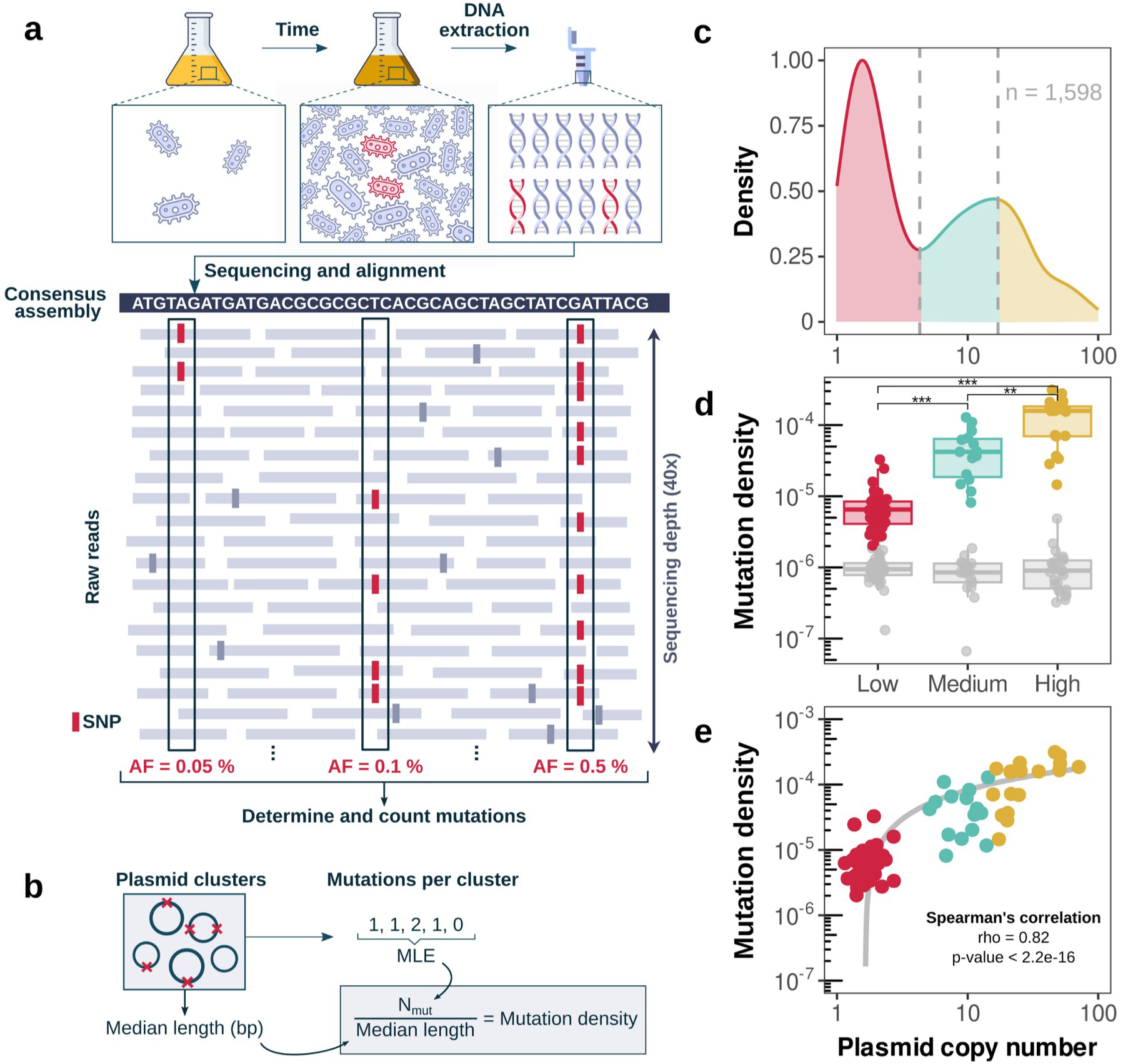
Mutation density determination using whole-genome sequencing data. **a)** This panel illustrates the method used to identify low-frequency mutations in whole-genome sequencing assays. First, cultures were grown to extract the genomic DNA. This growth time is sufficient to accumulate mutations within the culture population. Once the DNA was extracted and sequenced, we used the available reads in the NCBI database to search for these mutations. Reads are mapped against their respective assemblies to detect mutations within the “population” of reads, with a minimum allele frequency (AF) of 5% and a maximum of 50%. **b)** Schematic representation of how mutation density was estimated using sequencing data. For each plasmid cluster, the total number of detected mutations was used to determine the realized number of mutations per cluster (Nmut) using a maximum likelihood estimation (MLE) algorithm. Mutation density was then calculated by normalizing the number of realized mutations by the within-cluster median plasmid length. **c)** Distribution of PCN in E. coli plasmids (n = 1,598). Colors indicate thresholds used to classify plasmids as low, medium, or high PCN, based on the antimode (4.3 copies) and the second mode of the PCN distribution (17.06 copies). **d)** Plasmid mutation density (mutations per bp) by plasmid cluster (y-axis) for each condition of PCN: high (n = 27), medium (n = 17), and low (n = 40) (Kruskal-Wallis test (p < 10^−11^) followed by Wilcoxon rank sum test, low vs. medium, p < 10^−9^, effect size = 0.79; low vs. high, p < 10^−13^, effect size = 0.91; medium vs. high, p = 0.0018, effect size = 0.47, **Supplementary dataset 4**). The chromosomal mutation densities are shown in gray. **e)** Correlation between mutation density (mutations per bp) and PCN grouping by plasmid clusters (Spearman’s correlation, rho = 0.82 and p < 10^−16^). Gray line represents the Cannings model, where the constants c1 and c2 were adjusted to match the observed relationship between PCN and mutations per bp (**Supplementary dataset 3**). Only plasmids with fewer than 100 copies are shown.

We analyzed a previously published dataset comprising 723 *E. coli* genomes and 1,598 associated plasmids, for which complete assemblies, sequencing reads, and PCN estimates were available(Ramiro-Martínez et al. 2025). These plasmids belonged to 155 replicon types, 57 plasmid taxonomic units (PTUs), and 136 sequence-based clusters of highly similar plasmids (90% identity and k = 3 neighbours). We identified putative mutations by aligning sequencing reads to their corresponding consensus assemblies. To reduce potential false positives from sequencing errors, we implemented a strict filtering criterion: we retained only genomes with high overall coverage (≥30X) and considered only variants with allele frequencies of ≥5%(Garoña et al. 2021). Additionally, we excluded common Illumina-related artefacts by discarding SNPs unsupported by reads from both strands and variants located within homopolymeric regions(Deatherage and Barrick 2014) (see methods). This process resulted in the identification of 13,196 plasmidic and 192,322 chromosomal putative mutations.

In a classical Luria–Delbrück fluctuation experiment, parallel cultures are propagated, and the distribution of mutant counts across cultures is used to infer the realized number of mutation events(Luria and Delbrück 1943; Foster 1999; Barrick and Lenski 2013). Following a conceptually similar approach, we treated each plasmid within a sequence-based cluster as an independent replicate in a fluctuation assay and estimated the number of realized mutations per plasmid cluster using a maximum likelihood estimation algorithm(Zheng 2017). We then normalized mutation counts by median plasmid cluster length to obtain the mutation density, defined here as the number of within-cluster, high-confidence mutations per base pair (**Figure 4b**).

We first focused on host chromosomal mutation densities, and found that, as expected, they remained stable across PCN categories (**Figure 4d**, grey boxplots, Kruskal-Wallis test p = 0.57). Then, we classified plasmids into low, medium, and high copy number groups (**Figure 4c**) and compared their mutation density. High-copy plasmids exhibited significantly higher mutation densities than both medium-copy plasmids and low-copy plasmids (**Figure 4d**; Kruskal-Wallis test p < 10^−11^ followed by Wilcoxon rank-sum exact test, *p* < 0.0019, effect size > 0.28 for all pairwise comparisons; **Supplementary dataset 4**). Moreover, PCN and mutation density (**Figure 4e**) were strongly and positively correlated (Spearman’s rank correlation rho = 0.82, p < 10^−16^), even after normalising by chromosomal mutation density (**Supplementary Figure 13**, Spearman’s rank correlation rho = 0.80, p < 10^−16^). This relationship was well fitted by both a logarithmic model (R^2^ = 0.63, p < 10^−16^, **Supplementary dataset 2**) and the theoretical expectation derived from the Cannings model (residual sum-of-squares of 1.34×10^−7^ and one iteration with tolerance = 7.92×10^−9^). Of note, alternative methods for estimating plasmid and chromosomal mutation densities yielded qualitatively similar results (**Supplementary Figures 14-16**).

In agreement with the MA experiment, we observed a fivefold increase in mutation rates of low-copy plasmids compared with chromosomal rates. This consistent pattern suggests that plasmid replication may be inherently more error-prone, possibly due to the lower fidelity of the plasmid replication machinery or reduced access to DNA repair systems. Altogether, despite large variation in experimental conditions and other uncontrolled factors across datasets, our bioinformatic approach yielded a mutational signature that is highly compatible with our theoretical, computational and experimental observations.

## Discussion

The mechanisms of evolution do not operate independently; instead, highly intertwined and often opposing forces shape evolutionary trajectories. Plasmids, as autonomously replicating genetic elements, introduce an additional layer of complexity into bacterial evolution. On the one hand, plasmid replication typically ensures that multiple copies are present within each cell(Rodríguez-Beltrán et al. 2021; Ramiro-Martínez et al. 2025). This causes an increase in the availability of mutational targets relative to the chromosome, potentially accelerating the evolution of plasmid-encoded genes(San Millan et al. 2016; Hall et al. 2017; Sastre-Dominguez et al. 2024). On the other hand, because plasmid alleles must first reach fixation within individual cells before reaching fixation at the population level, plasmid-mediated adaptation can be significantly delayed(Ilhan et al. 2019; Santer and Uecker 2020; Garoña et al. 2021; Santer et al. 2022; Garoña et al. 2023; Dewan and Uecker 2025). These opposing forces have been studied separately, leading to controversial results that either support or refute the role of plasmids as catalysts of bacterial evolution(San Millan et al. 2016; Rodriguez-Beltran et al. 2018; Ilhan et al. 2019; Santer and Uecker 2020; Garoña et al. 2021; Santer et al. 2022; Garoña et al. 2023; Dewan and Uecker 2025; Pisera and Liu 2025).

We addressed this controversy by integrating theoretical, computational, experimental, and bioinformatic approaches. Using a Wright-Fisher-based model, we showed that plasmid evolution can be described as a Cannings process within the Kingman coalescent universality class, predicting a logarithmic relationship between PCN and mutation accumulation (**Figure 2**). Agent-based simulations supported these predictions, showing that plasmid genealogical tree lengths scale logarithmically with PCN. While different model formulations may affect the absolute number of mutations, the logarithmic relationship between PCN and mutation accumulation remains consistent. Notably, the fixed population size assumption central to classical population genetics is a natural feature of plasmid biology, making plasmids a compelling system for studying evolutionary dynamics through population genetics frameworks(Garoña et al. 2021; Hernandez-Beltran et al. 2022).

To empirically test these predictions, we performed an MA experiment using a plasmid with tunable PCN. Mutation rates markedly increased as PCN rose from ∼1 to 10 copies but seem to plateau beyond this point, even at 60 copies. These results were best explained by a logarithmic regression, which captured the decelerating relationship between PCN and mutation rates (**Figure 3**). Finally, we analyzed extensive sequencing data as independent fluctuation assays and showed that plasmids with higher PCNs tend to accumulate more detectable mutations per base pair (**Figure 4**). Together, these results indicate that plasmid evolvability scales logarithmically with PCN beyond laboratory conditions, with mutation rates plateauing around the natural boundary separating high- and low-copy plasmids (i.e., ∼6 copies per cell(Ramiro-Martínez et al. 2025)).

But what is the biological meaning of this logarithmic relationship? Population genetics offers an intuitive explanation: during severe bottlenecks and over many bacterial generations, most plasmid mutations are lost due to segregational drift, reducing their contribution to final plasmid diversity. However, over shorter time scales, the increase in mutational supply surpasses segregational drift, leading to a rise in the mutation rate associated with plasmids.

Consequently, under our study conditions, most of the plasmid mutations we detected likely arose shortly before sampling. This has broader implications for evolutionary analyses, as the rapid and recent accumulation of mutations in HCP can potentially complicate efforts to infer accurate evolutionary trajectories from endpoint data.

A crucial element of our theoretical, computational, and experimental (but not bioinformatic) analyses is using a single-cell bottleneck and the lack of selective forces. This maximizes segregational drift and minimizes the effective population size of plasmid lineages. Theory predicts that under less stringent bottlenecks, the effective mutation rate should be higher due to reduced effect of drift, suggesting that our estimates likely represent a lower bound. Future work applying similar approaches across a range of bottleneck sizes will help estimate effective mutation rates for plasmids in diverse ecological settings.

Beyond plasmids, our findings likely extend to any genetic element present in multiple copies and subject to random segregation during cell division, such as plastid and mitochondrial DNA (mtDNA). For example, human mitochondria contain thousands of mtDNA copies and undergo significant bottlenecks during maternal transmission(Árnadóttir et al. 2024). Vertebrate mtDNA exhibits a mutation rate approximately 20 times higher than nuclear DNA(Allio et al. 2017), a pattern that aligns with our results and underscores the robustness of the relationship between copy number and mutation rate across different genetic systems.

Collectively, our findings, along with other studies(San Millan et al. 2016; Rodriguez-Beltran et al. 2018; Zhang et al. 2019), highlight that plasmids are key drivers of bacterial evolution, extending beyond their role as gene delivery platforms. By functioning as autonomous genetic elements with increased mutation rates, plasmids can reduce the burden they cause to their host and potentiate the evolution of critical bacterial traits such as antibiotic resistance, virulence, and metabolism, underscoring their role as the evolutionary powerhouses of the bacterial cell.

## Materials and methods

### Bacterial strains, plasmids, and growth conditions

The strains used in this study were *E. coli* EPI300 and a hypermutator derivative of the same strain. To construct the hypermutator derivative strain, the *mutS* gene was replaced by a kanamycin resistance gene flanked by FRT sites, following the protocol described in ref. (Datsenko and Wanner 2000) with minor modifications. Briefly, the thermosensitive pKOBEG plasmid(Chaveroche 2000) carrying the lambda red system was introduced into the EPI300 strain. Then, a purified PCR product (primers MutS_F/R) amplified from the Δ*mutS::kan* KEIO mutant(Chaveroche 2000) was electroporated on arabinose-induced EPI300 cells. This PCR fragment contained the kanamycin resistance (*kan*) gene flanked by FRT sites, flanked by two ∼50 bp homology arms. After recombination, the pKOBEG plasmid was cured by incubating at 37°C. The thermosensitive pCP20 plasmid carrying site-specific FLP recombinase was then introduced to remove the kanamycin resistance, and subsequently cured by incubating at 42°C. Excision of the *kan* resistance gene was verified by PCR (primers K1/K2 and mutS_F/R), yielding the EPI300 Δ*mutS* strain. The hypermutator status of this strain was verified by performing fluctuation assays to rifampicin resistance.

EPI300 Δ*mutS* was transformed by heat shock with the pTA44 plasmid(10.3 kb)(Aakvik et al. 2009) as a tunable copy number plasmid. pTA44 carries two origins of replication: *ori2* and *oriV*(Aakvik et al., 2009). The *ori2* is a single-copy replication origin, while replication from *oriV* is proportional to the amount of TrfA initiation protein. The *trfA* gene, encoded in the host *E. coli* EPI300 Δ*mutS* chromosome, is controlled by the P_BAD_ promoter, which allows fine-tuning plasmid replication initiation by simply adding the inducer L-arabinose(Aakvik et al. 2009). Confirmation of the plasmid transformation was verified by PCR (repE_pTA44_F/R).

### Growth curves

Single colonies from each bacterial population were inoculated into LB starter cultures and incubated at 37°C for 16 hours with shaking at 225 rpm (three biological replicates). Each culture was then diluted 1:2000 in LB medium supplemented with kanamycin and the appropriate concentration of arabinose, and 200 μL of the diluted culture was transferred to a 96-well microtiter plate. The plates were incubated at 37°C with orbital shaking for 24 hours, and the optical density (OD) at 600 nm was measured every 10 minutes using a Synergy HTX plate reader (BioTek). The area under the growth curve (AUC) was calculated using the ‘auc’ function from the ‘flux’ R package. AUC was chosen as the primary growth metric because it incorporates key growth parameters, including maximum growth rate, lag phase duration, and carrying capacity.

### Mutation accumulation experiment

Each MA line was originated from a single colony independently isolated from LB agar plates. Each day, each replicate line was streaked to yield isolated colonies on LB agar plates with kanamycin and the appropriate concentration of arabinose and then incubated at 37°C for 22 hours. This process was repeated for 30 consecutive days. To ensure random selection of the colony to be passaged, the furthest colony on the streak’s end was chosen.

Initially, 12 *mutS−* lines were established for each condition of PCN. However, some lines were excluded due to highly similar mutation patterns that likely resulted from cross-contamination during the experiment. Specifically, the excluded lines were p0B1 and p0B3 from the 0% arabinose condition, p5A2, p5B1, and p5C4 from the 0.0005% arabinose condition, and p50A5 from the 0.5% arabinose condition. Additionally, one sample (p5B3) was removed due to a *trfA* missense mutation that caused a decline in PCN (**Supplementary Figure 12**). Consequently, ten lines were retained for the 0% arabinose condition, eight lines for the 0.0005% condition, and eleven lines for the 0.05% condition.

During the MA experiment, a random colony from each of the 12 lines was resuspended in 100 μL of MilliQ water daily to be stored at -20°C. Before storage, 20 μL of suspension was transferred to 180 μL of saline buffer, serially diluted, and replated to count viable cells and estimate the number of elapsed generations using the following formula: 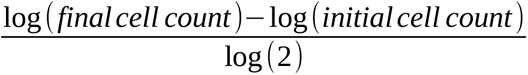. The stored samples were also used to estimate PCN every ten days during the experiment, as described below.

### Relative copy number determination

To determine relative PCN of the pTA44 plasmid, qPCR assays were performed using an Applied Biosystems 7300 Real-Time PCR System and NZYSupreme qPCR Green Master Mix (NZYTech), following the protocol described in ref.(San Millan et al. 2015). We developed a specific primers for pTA44 (repE_F: TCGGATTGACCTCTGCGGAAGC, repE_R: GCCTTTTTCATCGCCGGCATCC, amplicon size: 113 bp, efficiency: 99.09%, R^2^ = 0.999) and we used a previously described primer set for the dxs chromosomal gene (dxs_F: CGAGAAACTGGCGATCCTTA, dxs_R: CTTCATCAAGCGGTTTCACA, amplicon size: 113 bp, efficiency: 101.7%, R^2^ = 0.998)(Lee et al. 2006). Single colonies were resuspended in MilliQ water, heated at 98°C for 10 minutes, and directly used in qPCR reactions as template DNA(Škulj et al. 2008). Reaction efficiencies were determined from standard curves generated using five 8-fold serial dilutions of the template DNA, each tested in duplicate. qPCR cycling conditions included an initial polymerase activation step at 95°C for 2 minutes, followed by 40 cycles of denaturation at 95°C for 5 seconds and annealing/extension at 60°C for 30 seconds. Plasmid copy number was calculated using the following formula: PCN = [(1 + E_c_)^CTc^/(1 + E_p_)^CTp^] × (S_c_/S_p_), where E_c_ and E_p_ are the efficiencies of the chromosomal and plasmid qPCR, CTc and CTp are the threshold cycles of chromosomal and plasmid reactions, and S_c_ and S_p_ the size (bp) of the chromosomal and plasmid amplicons(San Millan et al. 2015).

Although qPCR is the most widely used method to estimate PCN, we validated these results using whole-genome sequencing data. Specifically, we calculated the PCN of each sample as the ratio of plasmid contig coverage to chromosomal coverage. Comparison of PCN values obtained by qPCR on the final day of the evolution experiment with those estimated from sequencing data revealed a strong and statistically significant correlation between the two methods (Pearson’s product-moment correlation, cor = 0.83, p = 9.7×10^−9^, **Supplementary Figure 17**).

### Whole Genome Sequencing and analysis of MA experiment

Thirty-six evolved populations of the hypermutator strain carrying the plasmid pTA44, were sequenced using the Illumina platform at the Wellcome Trust Center for Genomic Sequencing (Oxford, UK). Genomic DNA was extracted from 1 ml of resuspended cells using the Wizard Genomic DNA Purification Kit (Promega) following manufacturer’s instructions. All samples were quantified on a Qubit fluorometer (ThermoFisher) and a Nanodrop 2000c (ThermoFisher).

Raw Illumina sequencing reads were quality-filtered using TrimGalore v0.6.6 (https://github.com/FelixKrueger/TrimGalore), trimming low-quality bases from both ends with a minimum Phred quality score threshold of 20. Filtered reads were assembled de novo into contigs using SPAdes genome assembler v3.13.1 (https://github.com/ablab/spades) with default parameters. Assembled contigs were annotated using Prokka v1.14.5 (https://github.com/tseemann/prokka) to identify coding sequences and other genomic features. To identify plasmid-derived contigs for each strain, BLASTn v2.9.0+ was used to align all assembled sequences against the reference plasmid sequence.

To detect genetic variants in evolved populations relative to their respective ancestral strains, we used Breseq v0.36.1 (https://github.com/barricklab/breseq) (Deatherage and Barrick, 2014) in polymorphism mode, which enables the detection of both fixed and subpopulation (non-fixed) mutations. This approach was critical for capturing mutation rates across multicopy plasmids, where mutations may not be fixed in all copies. SNPs not supported by reads from both strands and variants located within homopolymeric regions were discarded, using the arguments *--polymorphism-coverage-both-strands 2* and *--polymorphism-reject-homopolymer-length 5*. The resulting mutation data were processed and summarized using custom scripts in R v4.1.0, and all statistical analyses were performed in R and RStudio (https://github.com/PaulaRamiro/Mutation_accumulation).

### Population Genetics Model of Plasmid Evolution

We described the dynamics of plasmid evolution using a Cannings model, a classical population genetics model that captures stochastic reproduction while maintaining a fixed population size. In our context, each generation corresponds to a bacterial reproduction event, and the population size at each generation is defined by the PCN, such that the key parameter N is equal to the number of plasmids per cell. During each reproduction event, plasmids have a 50% chance of being inherited by the daughter cell or remaining in the mother cell, independently of other plasmids. This results in a binomial distribution of plasmid counts in each new cell, with parameters *p*=1 /2 and *N*=*PCN* . A random binomial subset of plasmids, denoted as *B*∼ *Binomial* ( *N ,* 1/ 2), moves to the daughter cell, while the remaining *N* −*B* plasmids are replicated independently, each copying one of the *B* ancestors.

Mutations can occur during plasmid replication, with each mutation event occurring with a low probability. To capture the evolutionary dynamics of plasmid lineages, we treat plasmids as independent entities within a Cannings model with constant population size N (Cannings 1974), where at each generation, a random subset of plasmids is selected as ancestors for the next generation. Each plasmid in generation *g*+1descends from a uniformly chosen ancestor from the subset selected in generation *g*, and the remaining plasmids are replicated to restore copy number. The probability that two plasmids in a cell trace back to the same plasmid in the preceding cell is E[1/B], which is on the order of 1/N. Likewise, the probability that three plasmids share a common ancestor is E[1/B^2^], scaling as 1/N^2^. According to Möhle’s criteria (Möhle 1998), this places the genealogical process of plasmid lineages within the universality class of the Kingman Coalescent (Möhle and Sagitov 2001). This result implies that, under neutral conditions, the expected total branch length of the genealogy increases logarithmically with PCN, leading to a sublinear scaling of mutation accumulation with increasing copy number.

### Simulation of Plasmid Mutation Accumulation

The simulation was implemented in Python using standard scientific computing libraries, including NumPy and Matplotlib, with additional functionality from Biopython for lineage tree construction. The model represents bacterial cells as objects containing plasmids, each tracked independently to capture processes such as mutation accumulation and segregational drift. Plasmid replication was modelled as a stochastic process until reaching the maximum PCN, and segregation during cell division was simulated by randomly partitioning plasmids between daughter cells with a probability of 1/ 2.

The serial transfer protocol involved propagating populations for 60 simulated days, each comprising 24 generations. At the end of each day, a single bacterial cell was randomly selected to seed the next population. This bottleneck amplified genetic drift and allowed the accumulation of mutations to be observed over multiple transfers. Mutation events were introduced probabilistically during plasmid replication at a rate of _1.8 *×*10_^−8^ per plasmid per generation, with each mutation logged alongside its generation of origin. For each PCN value, we performed simulations until at least 1000 replicates with one or more mutations were obtained.

Lineage trees were constructed at the end of simulations by tracing plasmid ancestries based on replication and segregation events. The total length of these trees was calculated as the cumulative distance from root to tips, serving as a measure of genealogical divergence. Mutations were analyzed by tracking their frequency and persistence across generations to quantify key dynamics: emergence (appearance of new mutations), fixation (mutations reaching 100% frequency in the population), and loss (disappearance of mutations due to segregational drift or genetic bottlenecks). The simulation generated output files containing plasmid composition per cell across generations and reconstructed lineage trees, which were used to study genealogical divergence and mutation dynamics. The implementation is available at https://github.com/ccg-esb/MAp.

### Mutation rate analysis in *Escherichia coli* genomes

To detect mutations in naturally occurring plasmids, we first identified and downloaded all available assemblies from *Escherichia coli* annotated as “Complete Genomes” in the NCBI database (n = 4,124) on 05/12/2023. SRA information was extracted using the sra-toolkit v2.11.3 (https://github.com/ncbi/sra-tools) with a custom pipeline (https://github.com/PaulaRamiro/Mutation_accumulation) and used to download available paired-end reads (corresponding to n = 787 assemblies). Then, reads were trimmed to a Phred score of 20 with Trim Galore v0.6.6 (https://github.com/FelixKrueger/TrimGalore) and mapped against their respective assemblies with Breseq v0.38.1 (https://github.com/barricklab/breseq) with the -p flag to analyze non-fixed mutations within the populations. PCN information was extracted from ref.(Ramiro-Martínez et al. 2025).

We classified plasmids using different methods. First, we typed plasmids into different incompatibility groups according to their replication mechanism(Shintani et al. 2015; Robertson and Nash 2018) using MOB-typer from mob_suite v3.1.8 (https://github.com/phac-nml/mob-suite)(Robertson and Nash 2018) with the flag *--multi* to type independent plasmids within samples. This method analyzes features within the DNA sequences responsible for plasmid replication, such as the genes that encode replication initiation proteins. This allows us to establish plasmid groups that share similar replication mechanisms, known as replicon types. Second, we used a classification scheme based on similarity across the whole plasmid genetic content with COPLA v1.0(Redondo-Salvo et al. 2021). Plasmids that share high homology (>70%) in more than 50% of their sequence are assigned to the same plasmid taxonomic unit (PTU)(Redondo-Salvo et al. 2020; Redondo-Salvo et al. 2021). Third, we classified plasmids into different sequence-based plasmid clusters. Plasmid nucleotide FASTA files were sketched with Mash(Ondov et al. 2016) (k = 21, sketch = 10,000) to obtain an all-against-all pair-wise distance matrix; a k-nearest-neighbour graph was then built by connecting each plasmid to its three closest neighbours when the Mash distance was ≤ 0.1, and the resulting network was partitioned with the Fast-Greedy community-detection algorithm implemented in igraph(Csardi and Nepusz 2006), assigning every plasmid to a unique cluster.

We treated each plasmid as a replicate within groups categorized by clusters, replicon types, and PTUs as if they were replicates in a classical Luria–Delbrück fluctuation assay. The realized number of mutations (N_mut_) per plasmid group and 95% CIs were calculated using the maximum likelihood estimator (MLE), applying the *newton.LD* and *confint.LD* functions of the rSalvador package v1.9 for R(Zheng 2017). The p_0_ method was also used as a control to include the samples without mutations, with the *LD.p0.est* function. µ_eff_ for both chromosomes and plasmids was then calculated by dividing µ by the total length of the replicon in base pairs. This led to the estimation of µ_eff_ for 723 genomes and 1,598 plasmids.

## Competing interests

The authors declare no competing interests.

## Materials & Correspondence

Correspondence and requests for materials should be addressed to Rafael Peña-Miller and Jerónimo Rodríguez-Beltrán.

## Acknowledgements

We thank Álvaro San Millán, Javier DelaFuente, Alfonso Santos-López, and João Gama for their helpful suggestions. Work in the evodynamics lab (https://evodynamicslab.com/) is supported by project no. PI21/01363, funded by the Carlos III Health Institute (ISCIII) and co-funded by the European Union; CIBER -Consorcio Centro de Investigación Biomédica en Red-(CIBERINFEC) CB21/13/00084; Convocatoria SEIMC-FUNDACIÓN SORIA MELGUIZO de Investigación 2021; and funded by the European Union (ERC, HorizonGT, 101077809). Views and opinions expressed are, however, those of the author(s) only and do not necessarily reflect those of the European Union or the European Research Council Executive Agency. Neither the European Union nor the granting authority can be held responsible for them. PRM is a recipient of a predoctoral PFIS grant (grant no. FI22/00265) from the Carlos III Health Institute (ISCIII), through the Recovery, Transformation and Resilience Plan and Next Generation EU from the European Union. RPM received support from DGAPA-UNAM (PASPA-2024-III). JR-B acknowledges support by a Miguel Servet contract from the Carlos III Health Institute (ISCIII) (grant no. CP20/00154), co-founded by the European Social Fund, ‘Investing in your future’. We thank the Oxford Genomics Centre at the Wellcome Centre for Human Genetics (funded by Wellcome Trust grant reference 203141/Z/16/Z) for the generation and initial processing of sequencing data.

## Notes

### Competing Interest Statement

The authors have declared no competing interest.

