## Supplementary Information for "Plasmids promote bacterial evolution through a copy number-driven increase in mutation rate"

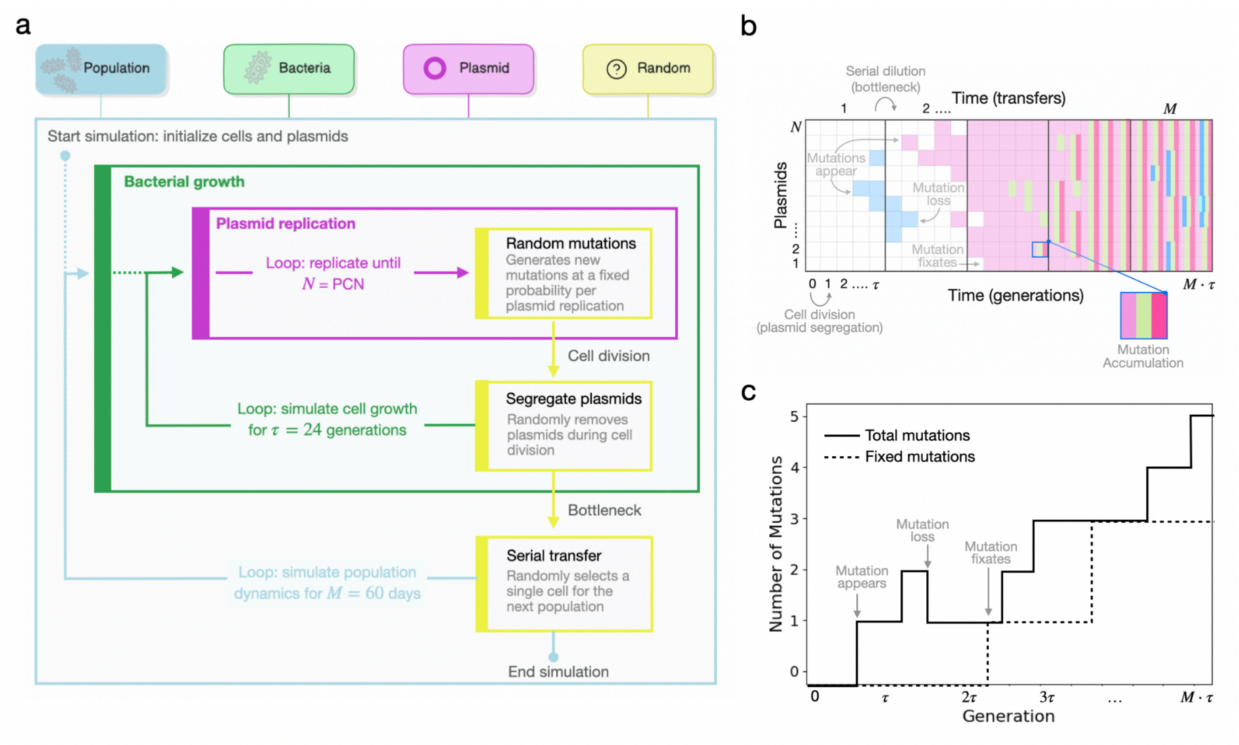


**Supplementary Figure 1. Schematic of the simulation workflow of the agent-based model.**

(a) The simulation begins by initializing a population of cells, each containing a defined copy number of mutation-free plasmids. During each daily cycle, plasmid replication is simulated by randomly selecting plasmids for replication until reaching a maximum copy number per cell. Mutations are introduced probabilistically during plasmid replication. After reaching the maximum plasmid number, with plasmids randomly partitioned between daughter cells. This process is repeated for 24 generations to simulate daily bacterial growth. At the end of each 24-generation cycle, a bottleneck occurs: a single cell is randomly selected to found the next day's population. Serial transfers are repeated for 30 days, allowing the accumulation of mutations within plasmid populations. The model explicitly tracks plasmid replication, mutation, and retention over time. (b) Example output of the agent-based simulation. The x-axis represents time, shown in generations (bottom axis) and corresponding serial transfers (top axis). The y-axis shows the number of plasmids present in the population. Each horizontal box represents a single plasmid at a given point in time. The color of each box encodes the mutation(s) carried by that plasmid; plasmids carrying multiple mutations are subdivided into segments, with each segment representing a distinct mutation. Arrows illustrate key events: plasmid segregation during cell division, the daily bottleneck caused by serial dilution (**τ** generations), and the appearance, loss, or fixation of mutations. This visualization allows tracing the fate of individual mutations and lineages over time. (c) Accumulated mutations over time in a single simulated lineage. The x-axis represents generations and the y-axis indicates the number of mutations present in the cell. The solid line shows the total number of mutations across all plasmids carried by the cell at each generation. The dashed line indicates the number of fixed mutations, defined as mutations present in every plasmid within the same cell.

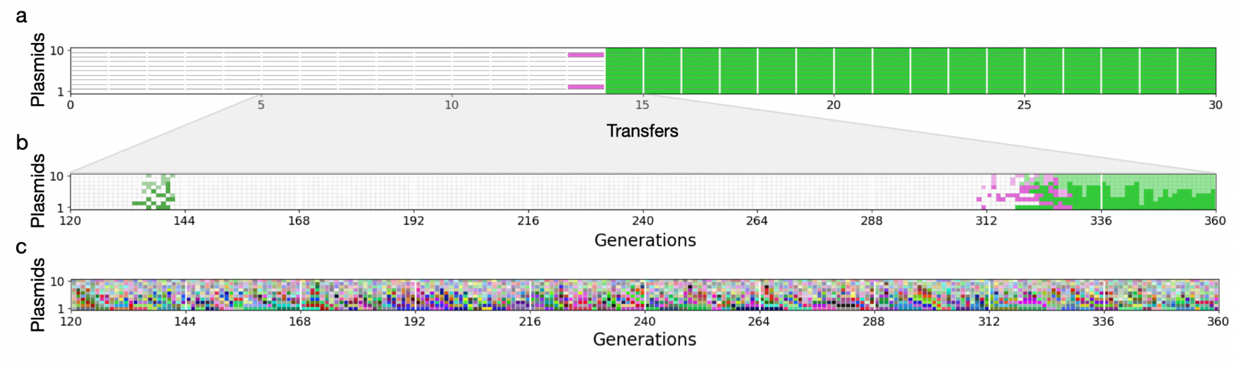


**Supplementary Figure 2. Representative simulation using the computational model (PCN = 10).**

Illustration of a stochastic simulation of plasmid evolution within a single cell with a PCN of 10 and a mutation rate of $10^{-3}$ per replication, simulated over 30 daily transfers. The x-axis represents time, and the y-axis corresponds to individual plasmids. (a) Mutations accumulated at the end of each transfer (every 24 generations), with each mutation shown in a different color. (b) Mutations recorded at the end of each generation during a zoomed-in window (generations 120 to 360), highlighting the dynamics of mutation accumulation at finer resolution. (c) Mutations recorded at the end of each generation during a zoomed-in window (generations 120 to 360), highlighting the dynamics of mutation accumulation at finer resolution. In panels (b) and (c), plasmids inherited from the previous generation are displayed with full opacity, while newly replicated plasmids appear semi-transparent.

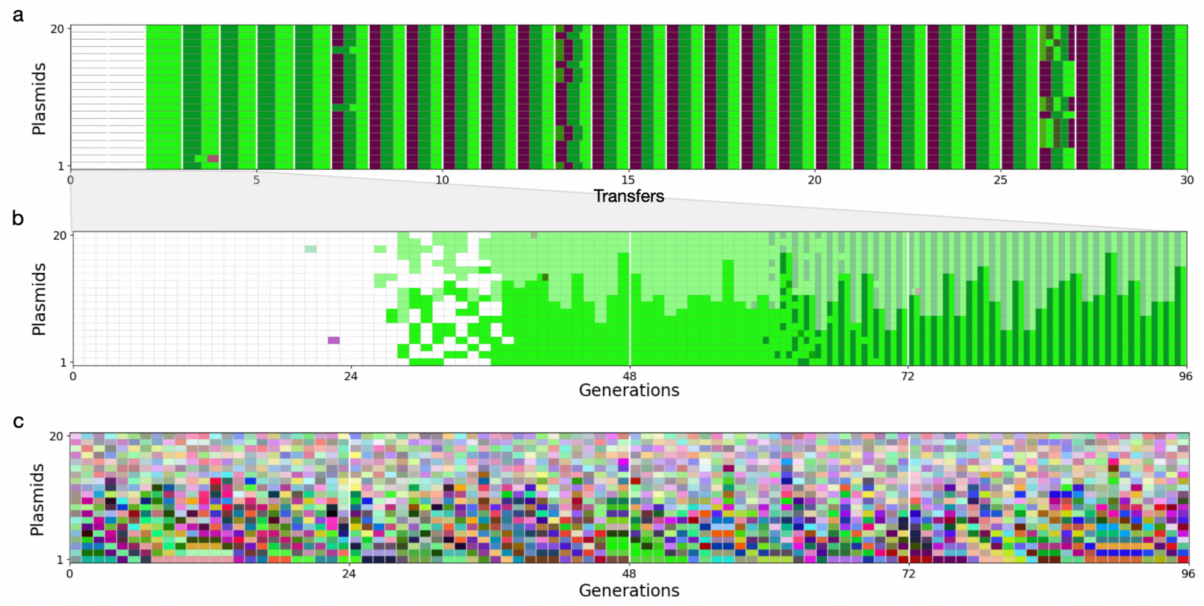


**Supplementary Figure 3. Representative simulation using the computational model (PCN = 20).**

Illustration of a stochastic simulation of plasmid evolution within a single cell with a PCN of 20 and a mutation rate of $10^{-3}$ per replication, simulated over 30 daily transfers. The x-axis represents time and the y-axis corresponds to individual plasmids. (a) Mutations accumulated at the end of each transfer (every 24 generations), with each mutation shown in a different color. (b) Mutations recorded at the end of each generation during a zoomed-in window (generations 0 to 96). Plasmids inherited from the previous generation are shown with full opacity, while newly replicated plasmids are represented by semi-transparent boxes. (c) Plasmid identity over time for the same zoomed-in window. Each plasmid is colored according to its unique identifier. Boxes with reduced opacity indicate newly replicated plasmids, whereas fully opaque ones were inherited from the previous generation.

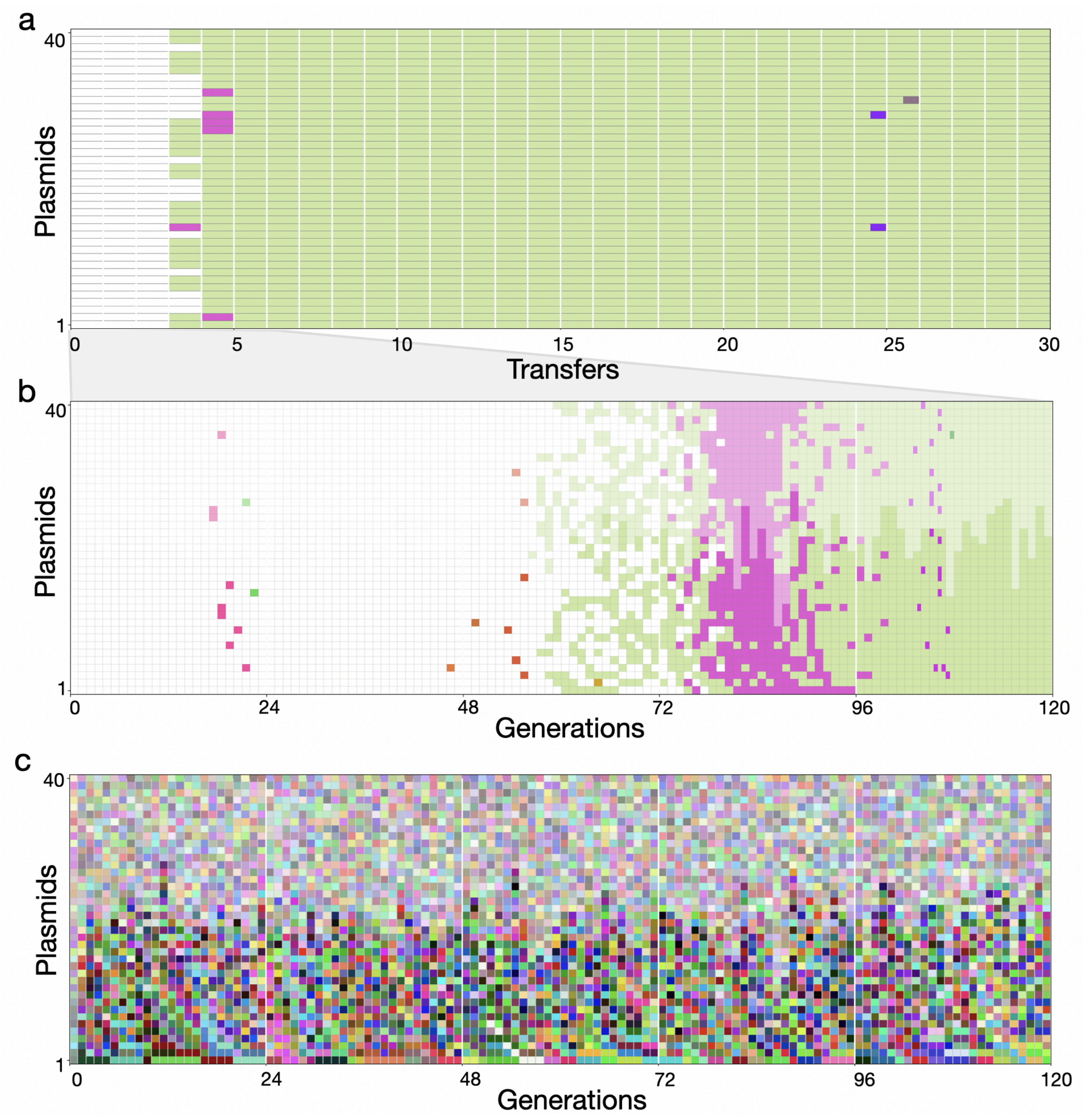


**Supplementary Figure 4. Representative simulation using the computational model (PCN = 40).**

Illustration of a stochastic simulation of plasmid evolution within a single cell with a PCN of 40 and a mutation rate of $10^{-3}$ per replication, simulated over 30 daily transfers. The x-axis represents time and the y-axis corresponds to individual plasmids. (a) Mutations accumulated at the end of each transfer (every 24 generations), with each mutation shown in a different color. (b) Mutations recorded after each generation during a zoomed-in window (generations 0 to 120). (c) Plasmid identity over time for the same zoomed-in window, where each plasmid is represented with a unique color. Boxes corresponding to plasmids inherited from the previous generation are shown with full opacity, while semi-transparent boxes indicate newly replicated plasmids.

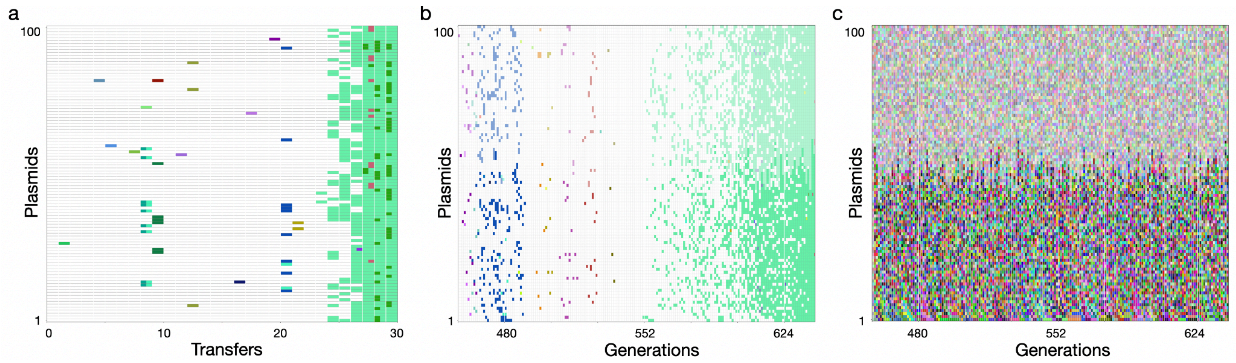


**Supplementary Figure 5. Representative simulation using the computational model (PCN = 100).**

Illustration of a stochastic simulation of plasmid evolution within a single cell with a PCN of 100 and a mutation rate of $10^{-3}$ per replication, simulated over 30 daily transfers. The x-axis represents time and the y-axis corresponds to individual plasmids. (a) Mutations accumulated at the end of each transfer (every 24 generations), with each mutation shown in a different color. (b) Mutations recorded at the end of each generation during a zoomed-in window (generations 456 to 640). (c) Plasmid identity during a zoomed-in window (generations 456 to 640), where each plasmid is represented with a different color.

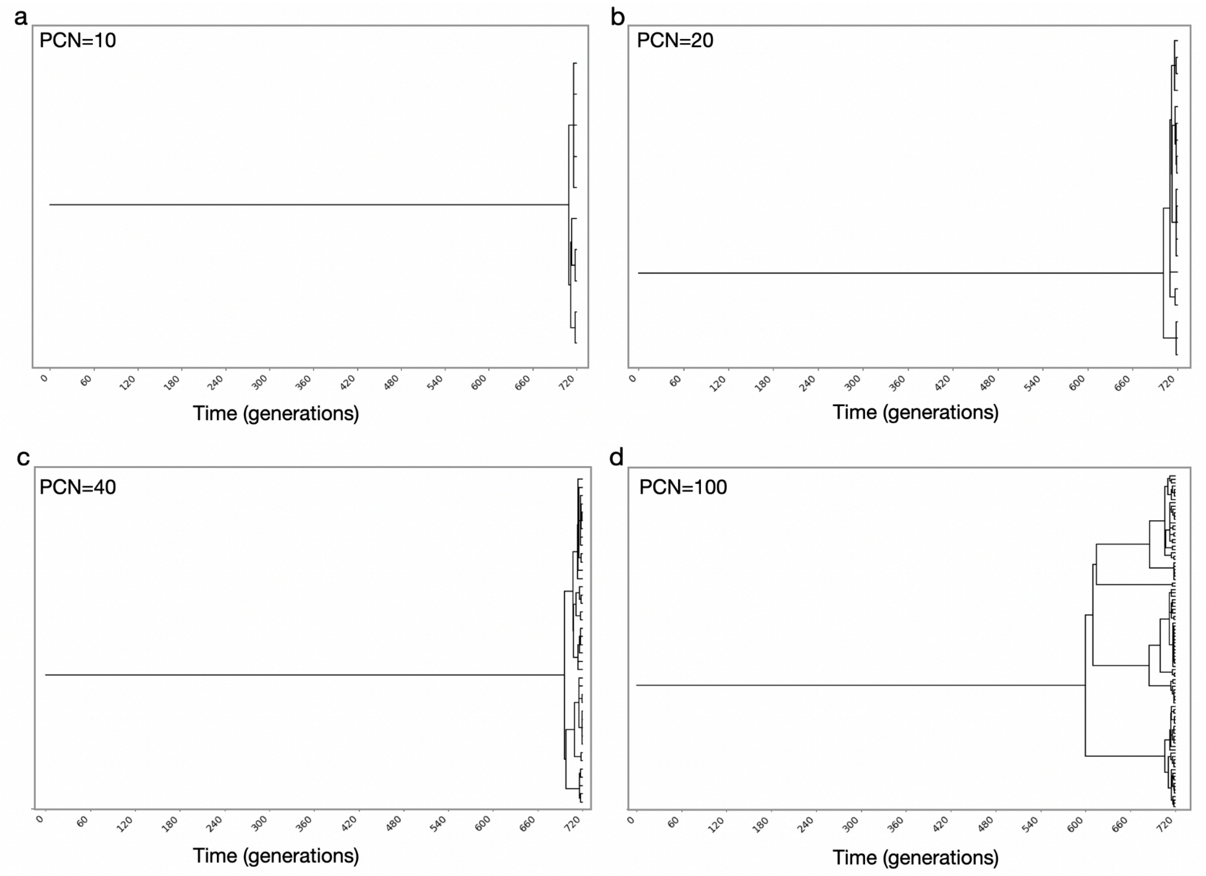


**Supplementary Figure 6. Plasmid lineage trees from single-cell simulations.**
Plasmid lineage trees corresponding to the single-cell simulations shown previously, for (a) PCN = 20, (b) PCN = 40, (c) PCN = 80, and (d) PCN = 100. Each tree displays a pattern of early coalescence followed by rapid branching, characteristic of clonally replicating plasmids that segregate independently during cell division.

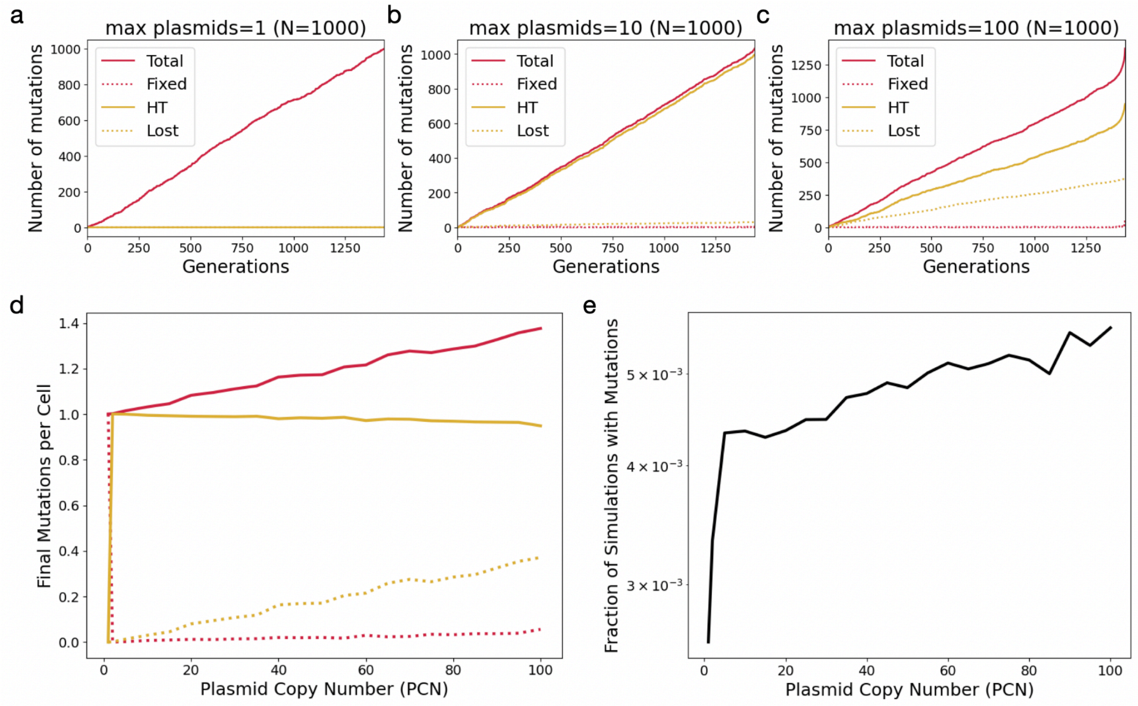


**Supplementary Figure 7. Effect of plasmid copy number on mutation accumulation dynamics.**

(a) Mutation accumulation over generations for PCN = 1 in 1000 stochastic simulations where at least one mutation was detected. (b) Mutation accumulation over generations for PCN = 10. (c) Mutation accumulation over generations for PCN = 100. Solid red lines indicate total mutations; dotted red lines show fixed mutations (present in all plasmids within a cell); solid yellow lines represent mutations present in a subset of plasmids (HT – heterozygous); and dotted yellow lines correspond to mutations lost through segregational drift. (d) Final number of mutations per cell for each mutation type as a function of PCN, using the same color scheme. (e) Fraction of all simulations that resulted in at least one mutation, plotted as a function of PCN (log scale).

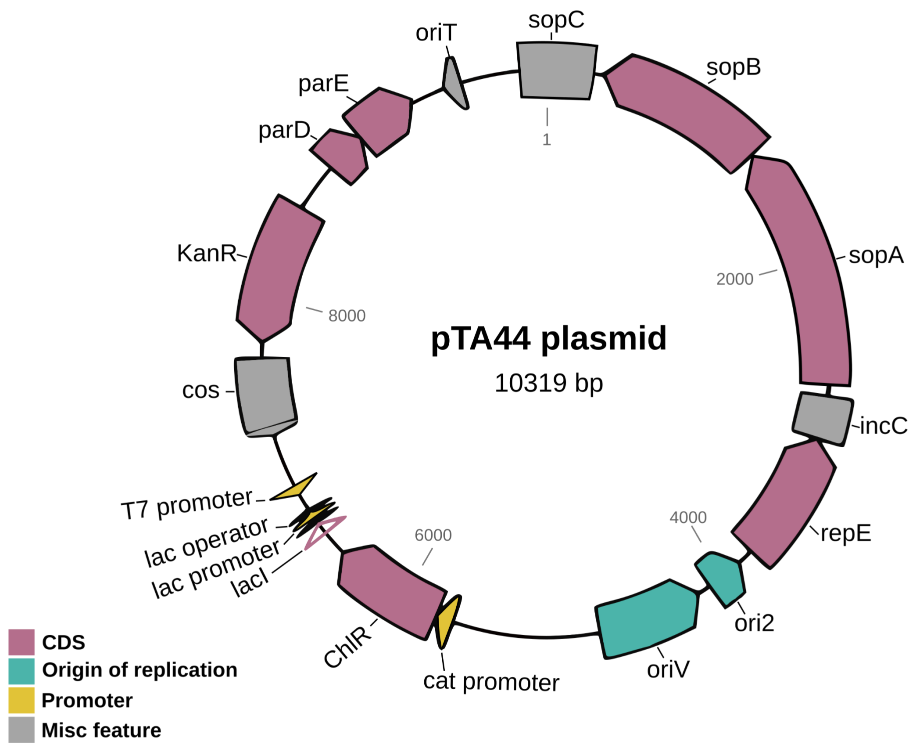


**Supplementary Figure 8. Genetic map of pTA44 plasmid.**

Coding sequences (CDS) are indicated by arrows. The two origins of replication (*ori2* and *oriV*) are marked in green. Promoters, antibiotic resistance genes, and other relevant genetic elements are annotated with distinct colors. Features were annotated and visualized using pLannotate (http://plannotate.barricklab.org/).

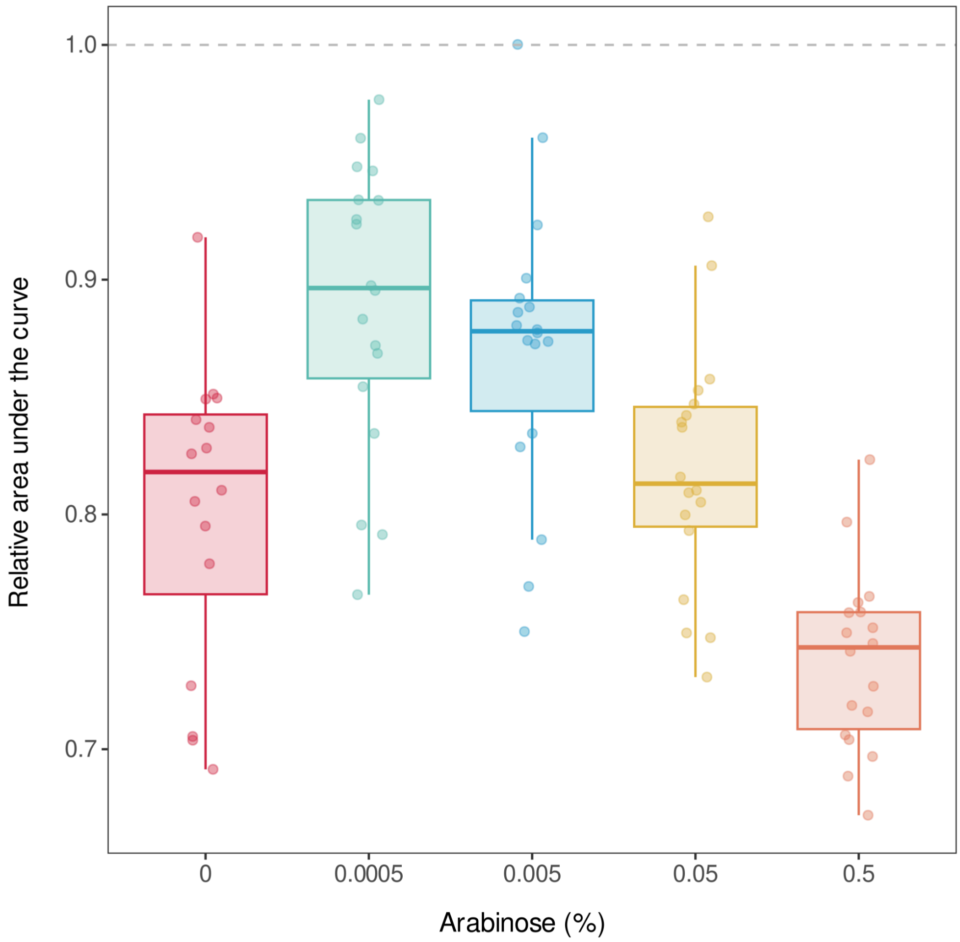


**Supplementary Figure 9. Fitness effects associated with PCN.**

Box plots illustrate the relative fitness of the E. coli EPI300 ΔmutS strain carrying the plasmid pTA44, compared to the plasmid-free strain at varying concentrations of arabinose (%). The horizontal lines within the boxes represent the median values, while the upper and lower hinges indicate the 75th and 25th percentiles, respectively. The whiskers extend to 1.5 times the interquartile range (IQR), and individual points represent independent technical replicates.

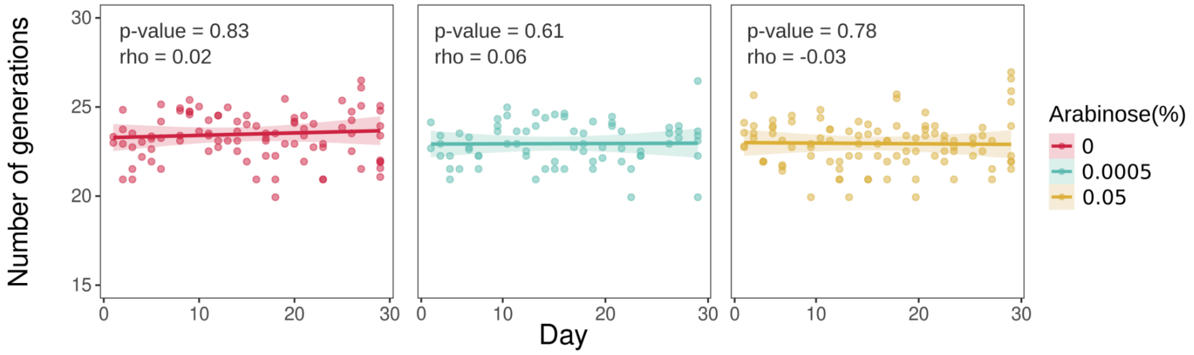


**Supplementary Figure 10. Number of generations during the experimental evolution.**

Scatter plots display the number of generations throughout the evolution experiment. Each data point represents an independent replicate. Spearman's rank correlation coefficient (rho) and associated p-values are indicated within each panel. Solid lines represent the best linear fits to the data, with shading indicating 95% confidence intervals around the fitted values.

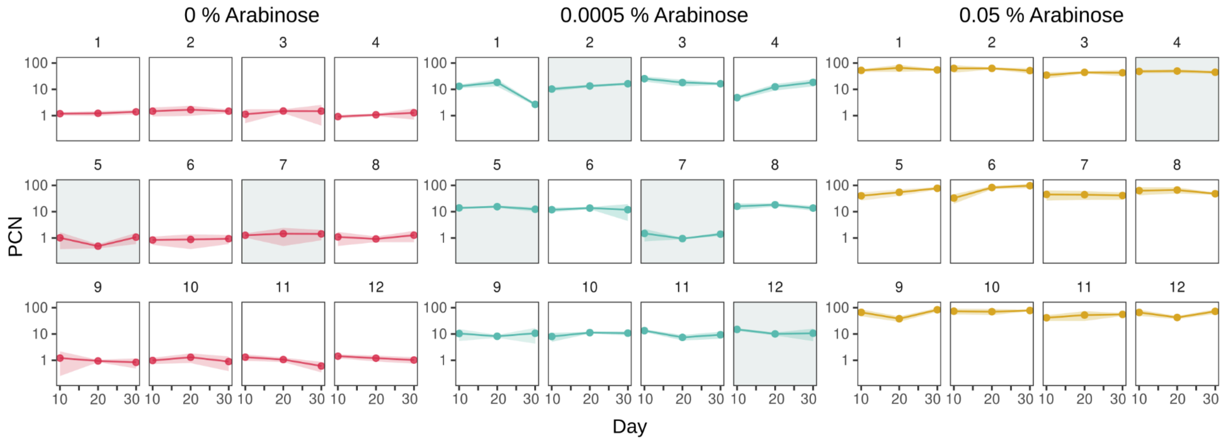


**Supplementary Figure 11. Plasmid copy number dynamics during the evolution experiment.**

Each panel represents the PCN dynamics of each population during the evolution experiment. The PCN is shown on a logarithmic scale on the y-axis, with time (in days) on the x-axis. Each panel represents an individual population. Points represent the median PCN of seven technical replicates. Points are connected with lines to aid visualization, with the shaded area indicating the interquartile range for each data point. Shaded panels indicate experimental lines that were excluded from further analyses, as detailed in the methods section.

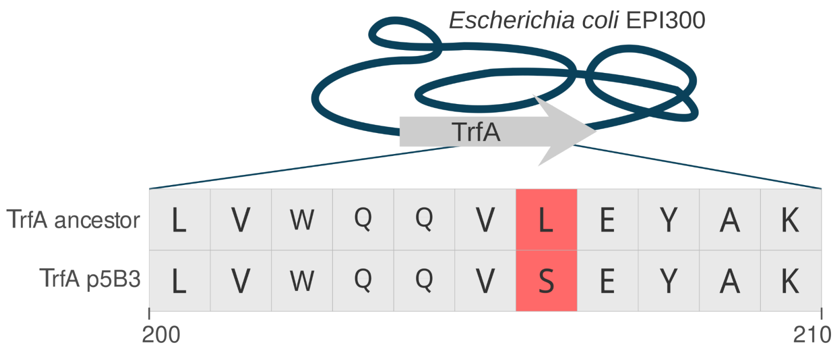


**Supplementary Figure 12. Mutation in the trfA gene on line 7 under 0.0005% arabinose condition.**

The figure illustrates an alignment covering amino acids from positions 200 to 210 of the ancestral TrfA protein (day 0) and the TrfA variant from line p5B3, which was evolved under 0.0005% arabinose. A missense mutation at position 206 leads to a substitution of leucine with serine (L206S), as highlighted in red. This mutation is associated with changes in plasmid copy number, likely due to a loss of TrfA functionality (refer to the main text for further details).

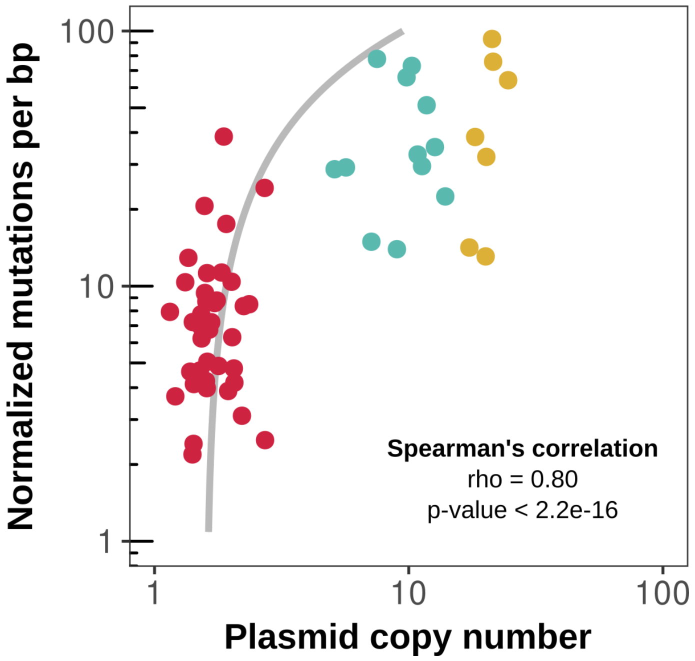


**Supplementary Figure 13. Correlation between mutations per bp normalized by chromosome mutation density and PCN.**

The mutation density of each plasmid (y-axis, measured in mutations per base pair) was normalized by the mutation density of its corresponding chromosome. Plasmids were categorized into three groups based on their copy number: low (red), medium (blue), and high (yellow) along the x-axis. The statistics for Spearman's correlation are displayed in the bottom right corner of the plot. The gray line represents the Cannings model, with constants c1 and c2 adjusted to align with the observed relationship between plasmid copy number and mutations per base pair (**Supplementary dataset 3**).

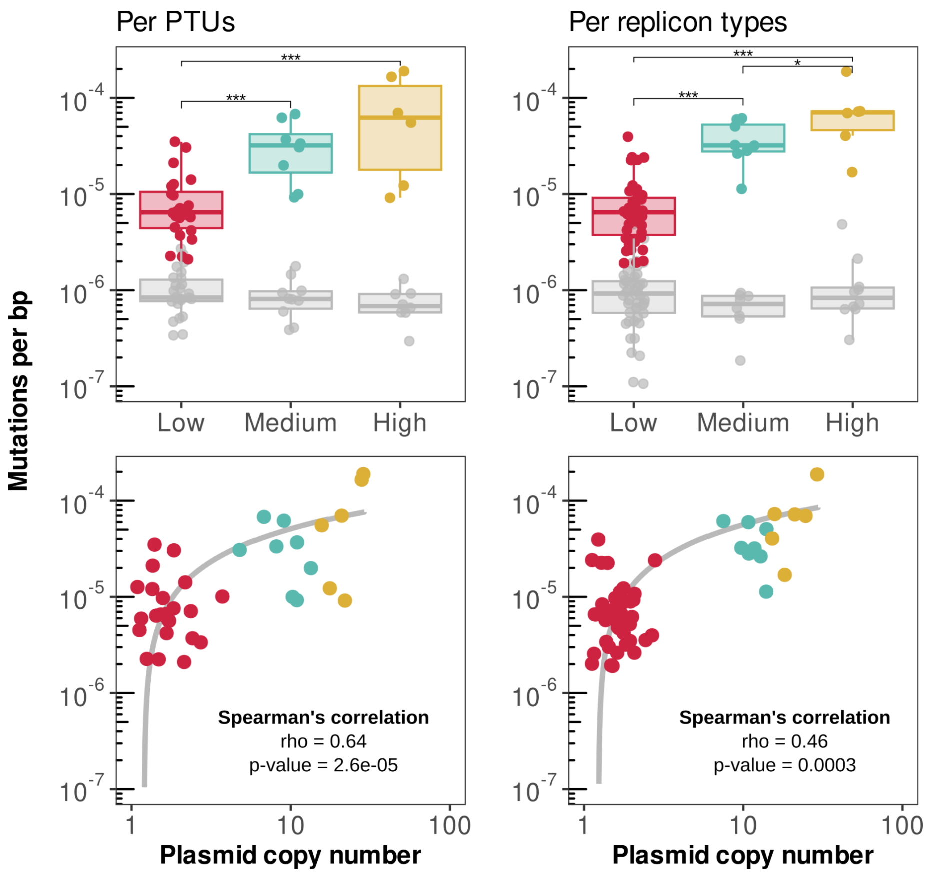


**Supplementary Figure 14. Mutation density determination by grouping the samples using different methods.**

Mutation density (y-axis, measured as mutations per base pair) were compared across plasmids grouped by plasmid taxonomic units (PTUs, left panels) or by replicon type (right panels). Each point represents a different plasmid group. Plasmids were classified into low (red), medium (blue), and high (yellow) plasmid copy number (x-axis).

Top panels indicate the differences between the groups (Kruskal-Wallis test ($p = 0.0002$) followed by Wilcoxon rank sum test, low vs. medium, $p = 0.00021$, effect size = 0.59; low vs. high, $p = 0.0015$, effect size = 0.56; medium vs. high, $p = 0.3450$, effect size =0.22 for PTUs; and Kruskal-Wallis test ($p < 10^{-7}$) followed by Wilcoxon rank sum test, low vs. medium, $p = 6.4x10^{-7}$, effect size = 0.68; low vs. high, $p = 6.4x10^{-7}$, effect size = 0.67; medium vs. high, $p = 0.04$, effect size = 0.48 for replicon types, **Supplementary dataset 4**). The chromosomal mutation densities are shown in gray for reference.

Bottom panels show correlations between PCN (log scale) and mutation density (Spearman's rank correlation rho = 0.64, $p < 10^{-5}$ for PTUs and rho = 0.46, $p = 0.0003$ for replicon types). Gray line represents the Cannings model, where the constants c1 and c2 were adjusted to match the observed relationship between PCN and mutations per bp (**Supplementary dataset 3**).

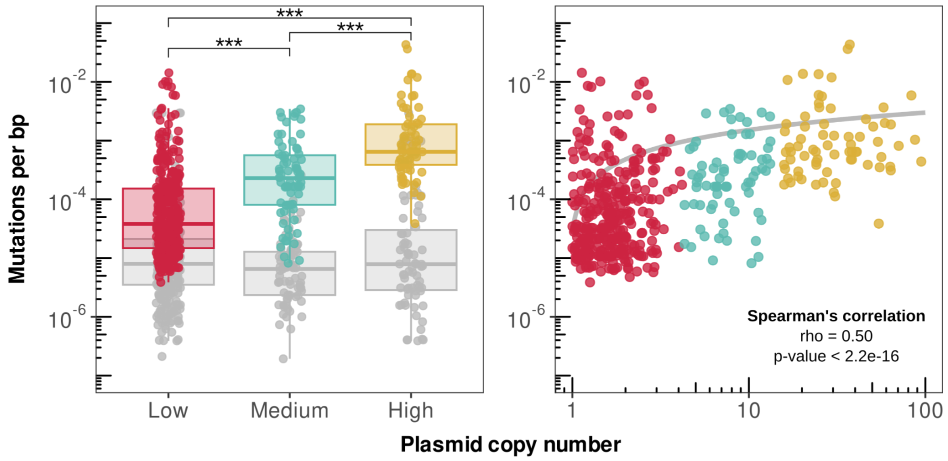


**Supplementary Figure 15. Mutations per bp for each plasmid.**

Mutation density (y-axis, measured as mutations per base pair) and plasmid copy number (x-axis) for each plasmid. Each plasmid is a different point (n= 1,598). Plasmids were classified into low (red), medium (blue), and high (yellow) plasmid copy number. Chromosomal mutation densities are shown in gray for reference. In the left panel, differences between the groups based on PCN are highlighted, (Kruskal-Wallis test ($p < 10^{-16}$) followed by Wilcoxon rank sum test, low vs. medium, $p = 7.2x10^{-10}$, effect size = 0.309; low vs. high, $p < 10^{-1}$, effect size = 0.555; medium vs. high, $p = 2,6x10^{-9}$, effect size = 0.478, **Supplementary dataset 4**). The right panel shows the correlation between PCN (log scale) and mutation density (Spearman's rank correlation rho = 0.50, $p < 10^{-16}$). Gray line represents the Cannings model, where the constants c1 and c2 were adjusted to match the observed relationship between PCN and mutations per bp (**Supplementary dataset 3**).

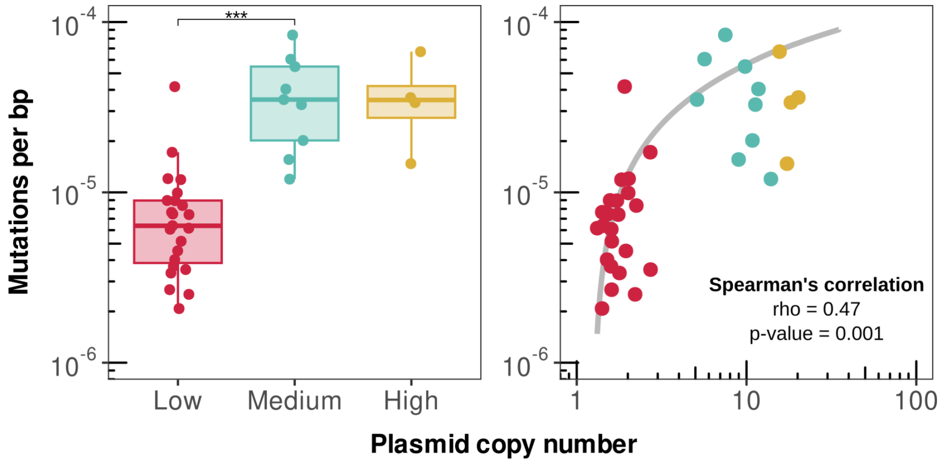


**Supplementary Figure 16. Mutation density with the $p_0$ method.**

Mutation density (y-axis, measured as mutations per base pair) and plasmid copy number (x-axis) for each plasmid cluster. Each point represents a different cluster. Plasmids were classified into low (red), medium (blue), and high (yellow) plasmid copy number. In the left panel, differences between the groups based on PCN are highlighted, (Kruskal-Wallis test ($p = 0.0003$) followed by Wilcoxon rank sum test, low vs. medium, $p = 2x10^{-6}$, effect size = 0.66; low vs. high, $p = 0.31$, effect size = 0.15; medium vs. high, $p = 0.4$, effect size = 0.13, **Supplementary dataset 4**). The right panel shows the correlation between PCN (log scale) and mutation density (Spearman's rank correlation rho = 0.47, $p = 0.001$). Gray line represents the Cannings model, where the constants c1 and c2 were adjusted to match the observed relationship between PCN and mutations per bp (**Supplementary dataset 3**).

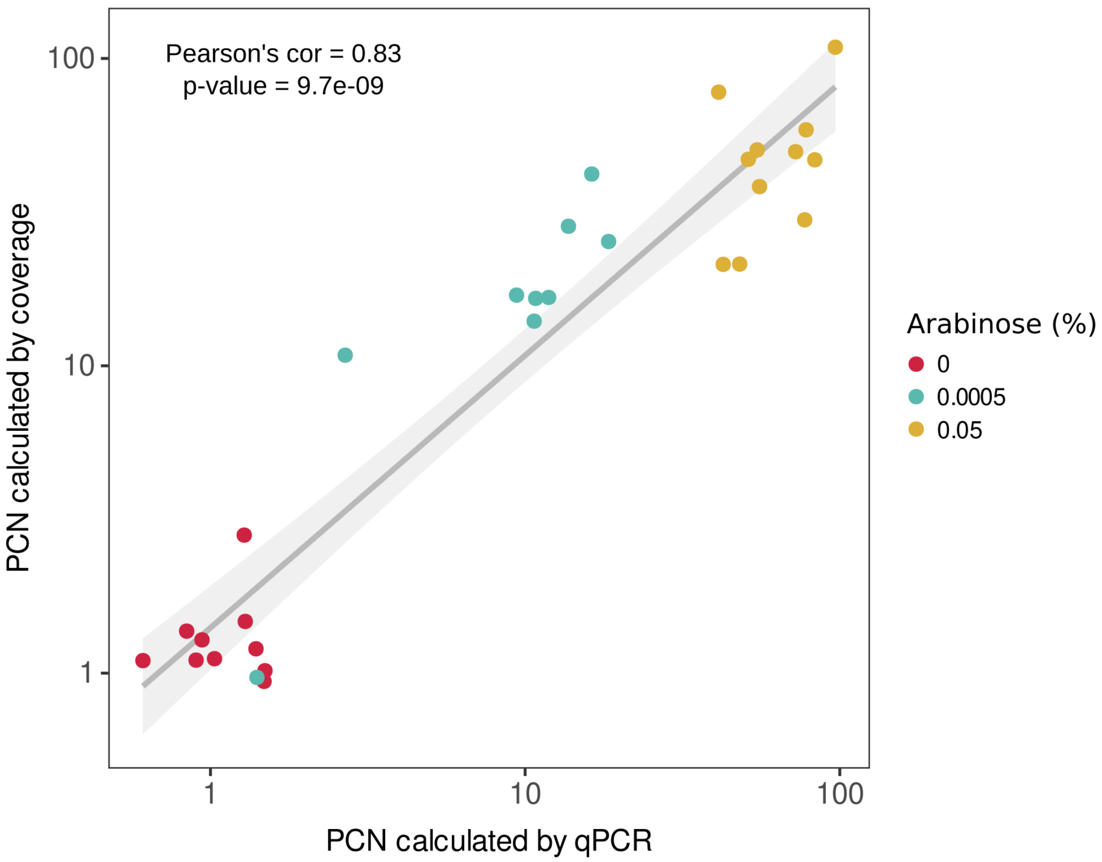


**Supplementary Figure 17. Correlation between PCN estimated by qPCR and sequence coverage.**

Scatter plot showing PCN values for individual samples calculated using qPCR (x-axis) and sequencing read coverage (y-axis), plotted on a log-log scale. Each point represents a single sample, colored according to the arabinose condition. Statistics of Pearson's correlation are indicated within the plot. The solid line represents the best linear fit, with the shaded area indicating the 95% confidence interval.
